# Patterns of putative gene loss suggest rampant developmental system drift in nematodes

**DOI:** 10.1101/627620

**Authors:** Gavin C. Woodruff

## Abstract

Gene loss often contributes to the evolution of adaptive traits. Conversely, null mutations frequently reveal no obvious phenotypic consequences. How pervasive is gene loss, what kinds of genes are dispensable, and what are the consequences of gene loss? The nematode *Caenorhabditis elegans* has long been at the forefront of genetic research, yet only recently have genomic resources become available to situate this species in its comparative phylogenetic and evolutionary context. Here, patterns of gene loss within *Caenorhabditis* are evaluated using 28 nematode genomes (most of them sequenced only in the past few years). Orthologous genes detected in every species except one were defined as being lost within that species. Putative functional roles of lost genes were determined using phenotypic information from *C. elegans* WormBase ontology terms as well as using existing *C. elegans* transcriptomic datasets. All species have lost multiple genes in a species-specific manner, with a genus-wide average of several dozen genes per species. Counterintuitively, nearly all species have lost genes that perform essential functions in *C. elegans* (an average of one third of the genes lost within a species). Retained genes reveal no differences from lost genes in *C. elegans* transcriptional abundance across all developmental stages when considering all 28 *Caenorhabitis* genomes. However, when considering only genomes in the subgeneric *Elegans* group, lost genes tend to have lower expression than retained genes. Taken together, these results suggest that the genetics of developmental processes are evolving rapidly despite a highly conserved adult morphology and cell lineage in this group, a phenomenon known as developmental system drift. These patterns highlight the importance of the comparative approach in interpreting findings in model systems genetics.

## Introduction

Gene loss is common and often has phenotypic consequences. Such losses, whether due to large-scale structural variation or to single nucleotide changes that render proteins non-functional, are typically associated with disease states that can represent profound public health challenges (Stankiewicz and Lupski 2010). However, gene loss also frequently underlies adaptive change (Albalat and Cañestro 2016). Such losses underlie changes in leaf morphology among Brassicaceae plant species (Vlad, et al. 2014), cold temperature resistance in flies (Greenberg, et al. 2003), self-incompatibility in *Arabidopsis* (Shimizu, et al. 2008), and pigmentation variation in multiple systems (Zufall and Rausher 2004; Protas, et al. 2006; Hoballah, et al. 2007). Similarly, selection can drive gene loss or genome size reduction in the context of experimental evolution as well (Nilsson, et al. 2005; Good, et al. 2017). The absence of a gene can even promote reproductive isolation and thereby play an important role in speciation (Bikard, et al. 2009; Ben-David, et al. 2017). Thus, gene loss must contribute to evolutionary change. What kinds of genes are dispensable, and how might they promote phenotypic divergence?

Conversely, although gene loss often has dramatic phenotypic consequences, a common outcome of gene loss is no observable phenotypic consequence at all. For instance, although the average human being is homozygous null for about twenty genes, most people do not have genetic disorders (MacArthur, et al. 2012). In addition, multiple large-scale knockout and knockdown screens for genetic function have unearthed thousands of genes with no obvious function in multiple organisms (Winzeler, et al. 1999; Kamath, et al. 2003; Dietzl, et al. 2007). Such observations are often explained by genetic redundancy, wherein multiple genes perform the same function, and therefore the loss of any one such gene is of little phenotypic consequence (Nowak, et al. 1997). However more recent studies have revealed that the fitness consequences of many gene knockdowns vary depending on genetic background (Dowell, et al. 2010; Chari and Dworkin 2013; Paaby, et al. 2015). Here such results could be explained by pervasive compensatory change and developmental system drift (True and Haag 2001), wherein dramatic differences in underlying developmental processes nonetheless promote similar phenotypes. Overall, then, the extent to which gene loss influences phenotypic change (or lack thereof) is not completely understood.

The first metazoan to have its genome sequenced, the nematode *C. elegans* has been a widely used model system for decades (Corsi, et al. 2015). In addition to its widespread use in developmental and molecular genetics, much is known about its genomic features (Gerstein, et al. 2010). Indeed, the WormBase database contains vast information for many of its ~20,000 protein-coding genes (Howe, et al. 2016). Despite this, the comparative and evolutionary genomic resources of *C. elegans* have been historically lacking compared to other widely studied model systems such as *Drosophila* (Consortium 2007; Huang, et al. 2014; Casillas and Barbadilla 2017). However, this has recently changed with the rapid discovery of dozens of *Caenorhabditis* species (Kiontke, et al. 2011; Ferrari, et al. 2017; Slos, et al. 2017; Stevens, et al. 2019) in tandem with the sequencing of multiple close relatives of *C. elegans* (Fierst, et al. 2015; Slos, et al. 2017; Kanzaki, et al. 2018; Ren, et al. 2018; Rödelsperger 2018; Yin, et al. 2018; Stevens, et al. 2019). Here, I combine the collective knowledge of *C. elegans* developmental genetics that resides in the WormBase database with patterns of gene loss observed across the genomic evolution of 28 *Caenorhabditis* species, finding that patterns of species-specific gene loss underlie vast developmental system drift in this genus. These patterns underscore the crucial role of genomic context in understanding gene function.

## Results

### *All* Caenorhabditis *species have lost multiple genes that perform essential functions in* C. elegans

To explore the extent and consequences of gene loss in *Caenorhabditis*, species-specific gene losses were inferred at two levels of phylogenetic scope (the whole genus and only the *Elegans* group, Figure 1). Briefly, after defining groups of orthologous proteins across 28 *Caenorhabditis* species (Emms and Kelly 2015), orthologous groups present in all species but one were called presumptive species-specific lost genes. Here, gene loss will be assumed to be equivalent to this type of species-specific absence, as opposed to many other possible patterns of repeated loss in multiple species; here I only examined patterns of species-specific loss.

**Figure 1.**
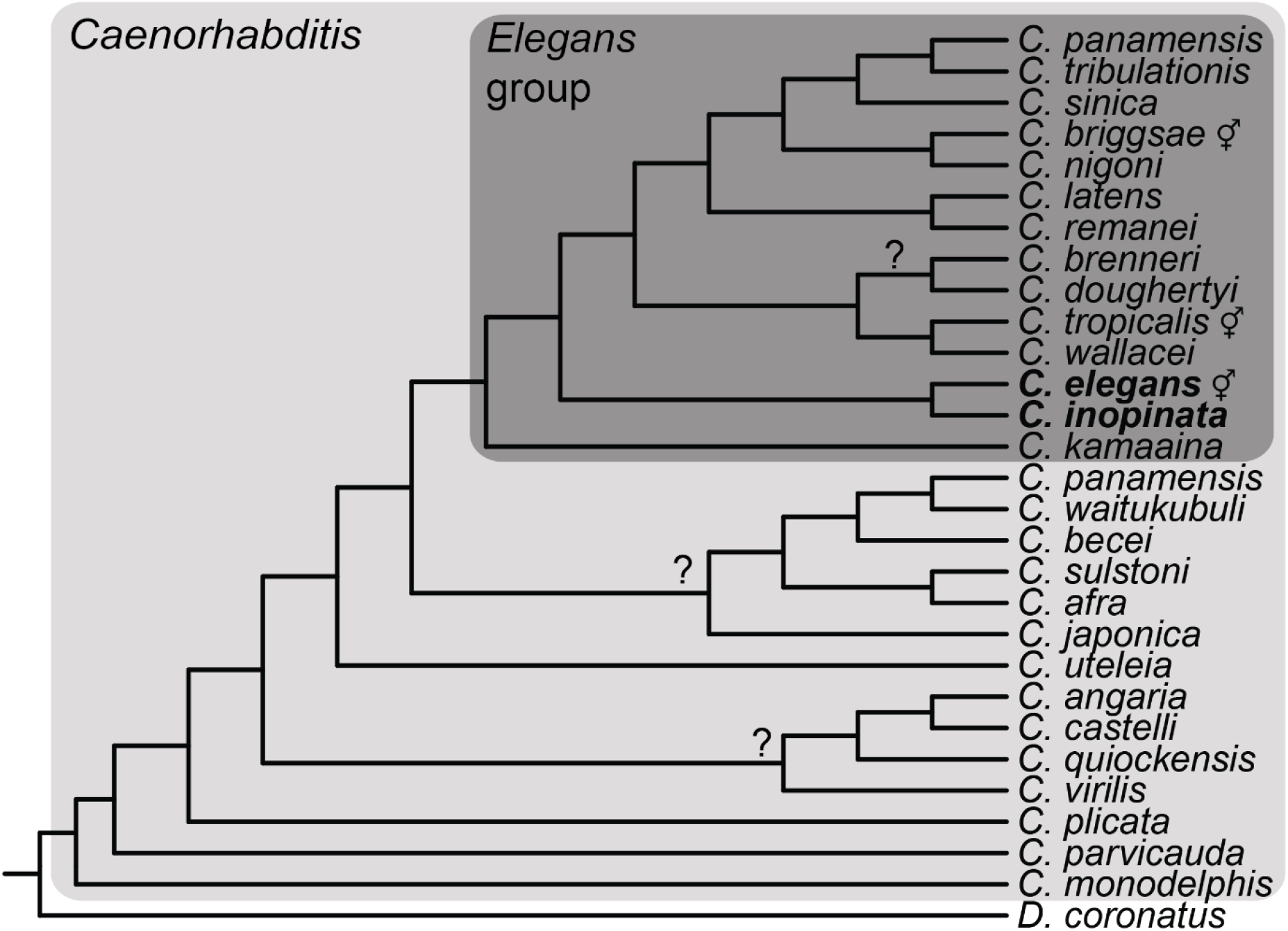
Cladogram showing taxa used and phylogenetic levels of analysis (species-specific loss across all *Caenorhabditis*; species-specific loss in the *Elegans* group; only lost in *C. inopinata*). It is important to note that the species included in this study do not constitute all available *Caenorhabditis* genomes nor known *Caenorhabditis* species (Kiontke, et al. 2011; Slos, et al. 2017). Throughout this manuscript, “all *Caenorhabditis*” refers to all *Caenorhabditis* species included here. The cladogram is based on the Bayesian phylogenetic analysis in Stevens et al. 2019 (Stevens, et al. 2019). Question marks represent nodes with low support or incongruence among methods of phylogenetic inference (Stevens, et al. 2019). ⚥, hermaphroditic species.

At both phylogenetic levels, all species exhibit multiple species-specific gene losses (Figure 2). As the number of shared orthologous groups declines as more species are included (Supplemental Figure 6), the number of gene losses per species is higher when considering the *Elegans* group (mean=96, median=40, range=11-556) than when considering the genus as a whole (mean=48, median=19, range=2-201). As the genome assemblies under consideration are at varying degrees of completeness and quality (Supplemental Figures 3-5), this may have influenced the number of inferred species-specific losses. However, gene loss is only significantly associated with genome completeness when considering the *Elegans* group (Supplemental Figure 8) and not the whole genus (Supplemental Figure 9). Furthermore, gene loss is not significantly correlated with scaffold number (Supplemental Figures 10-11) or N50 (Supplemental Figure 12-13). Thus, although genome quality may influence the inference of gene loss, and the results here may overestimate gene loss, most genome quality metrics are not correlated with gene loss. Moreover, there are a number of species with high quality reference genomes (*C. elegans*, *C. briggsae*, *C. tropicalis*, *C. nigoni*, *C. wallacei*, and *C. inopinata*), and all of these species exhibit species-specific gene loss (Figure 2). Indeed, in the case of *C. inopinata*, the degree of gene loss with respect to the *Elegans* group is high (169 lost orthologous groups), despite its completely assembled reference genome (seven, chromosome-level scaffolds) and high BUSCO completeness score (98.1%)(Kanzaki, et al. 2018). So although genome quality likely inflates the extent of gene loss in some species, patterns of species-specific gene loss are still detected even in high quality assemblies.

**Figure 2.**
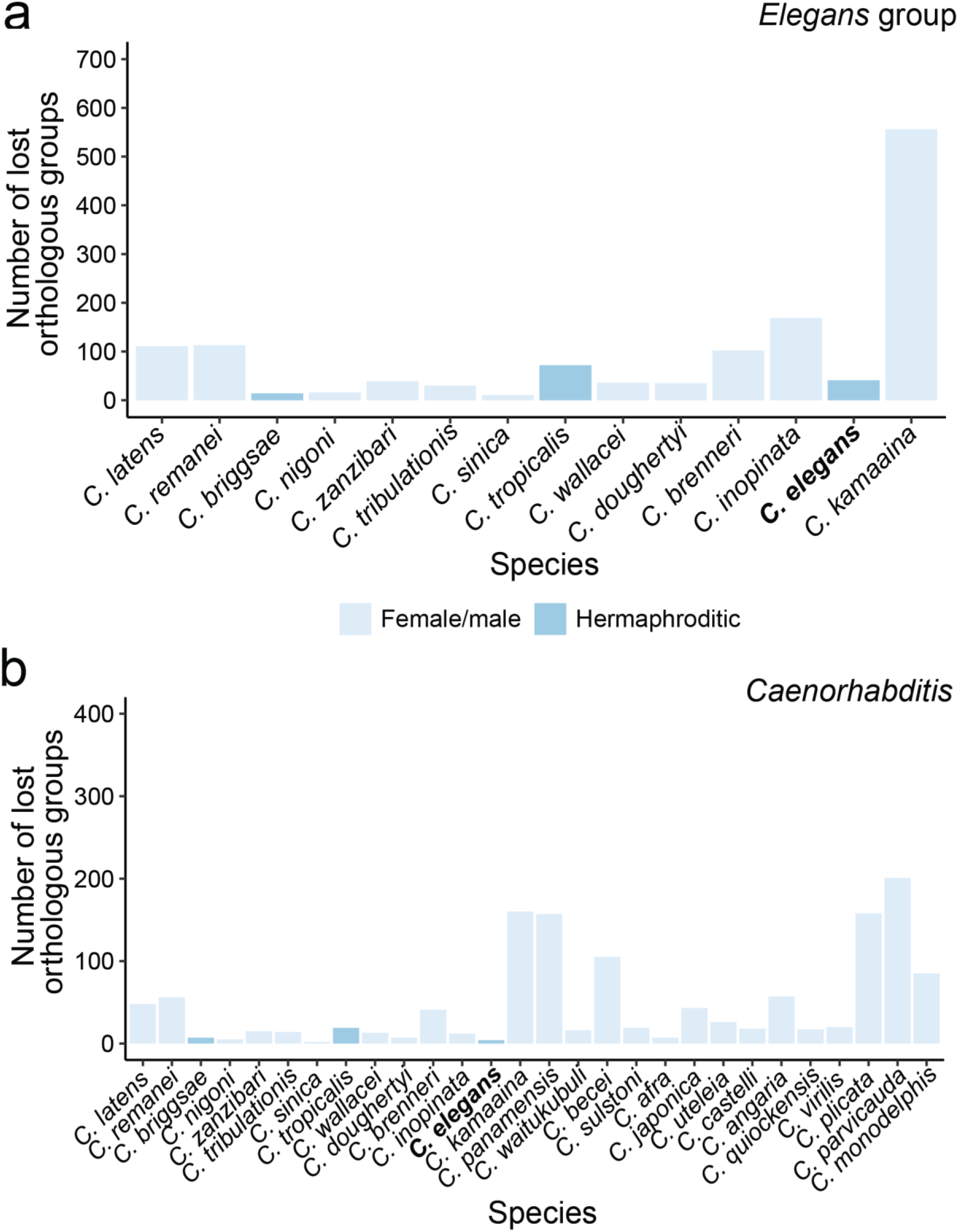
All species have uniquely lost multiple orthologous genes. Species-specific orthologous group losses when considering only the *Elegans* group (a) or all *Caenorhabditis* species (b). Bars are colored by the reproductive mode of the species. Species are roughly ordered by phylogenetic relatedness.

To understand their functional relevance, lost genes were paired with WormBase phenotype data ((Schindelman, et al. 2011); see methods). WormBase is a repository for biological knowledge of *C. elegans*, notable for housing various kinds of genomic data related to *C. elegans* and its relatives (Howe, et al. 2015). Among these are “phenotype” terms, which constitute a formal ontology used to describe phenotypes associated with genes (Schindelman, et al. 2011). More specifically, these describe biological phenotypes that arise upon some perturbation of a given gene, usually through mutation or RNAi knockdown (Schindelman, et al. 2011). There are at least 2,443 phenotype terms in WormBase, ranging from the straightforward (“embryonic lethal,” “no germ line”) to the esoteric (“loss of asymmetry AWC,” “nuclear fallout”). I paired all of the *C. elegans* gene constituents of lost orthologous groups in each species at both phylogenetic levels with their WormBase phenotype terms. Among genes with phenotypes (only about 42% of *C. elegans* protein-coding genes were found to have phenotypes in WormBase (8,514 out of 20,204)), I further classified them into two categories: essential and inessential. Essential phenotypes were defined by the presence of any of these words: “lethal,” “arrest,” “sterile,” or “dead,” and ultimately constituted 58 unique phenotype terms (see Supplemental Data for list of essential phenotypes). All other phenotypes were noted as inessential. It is important to note that there are multiple phenotypes here noted as inessential that are probably critical for survival and that these numbers of essential genes reported here are likely underestimates. Additionally, the phenotypes of genes lost only in *C. elegans* cannot be assessed because WormBase phenotypes are derived from studies of genes that are present in *C. elegans*.

At both phylogenetic levels considered, nearly all species have lost genes that are associated with both essential and inessential phenotypes (Figure 3). Among the *Elegans* group, 36% of the *C. elegans* genes associated with species-specific lost orthologous groups had phenotypes on average (range=23%-53%), and 20% had essential phenotypes (range=0%-36%; Figure 3a). Across all *Caenorhabditis*, 37% of the *C. elegans* genes associated with species-specific lost orthologous groups had phenotypes on average (range=17%-71%), and 23% had essential phenotypes (range=0%-50%; Figure 3b). Notably, *C. briggsae* has not lost any essential genes at both levels of phylogenetic consideration; its close relative *C. nigoni* has also lost no essential genes when considering the whole genus (although it has lost four essential genes when considering the *Elegans* group; Figure 3). All other species have lost genes that are needed for viability and fecundity in *C. elegans*. These patterns suggest that genetic functions among highly conserved processes (such as embryogenesis) are rapidly evolving in this group.

**Figure 3.**
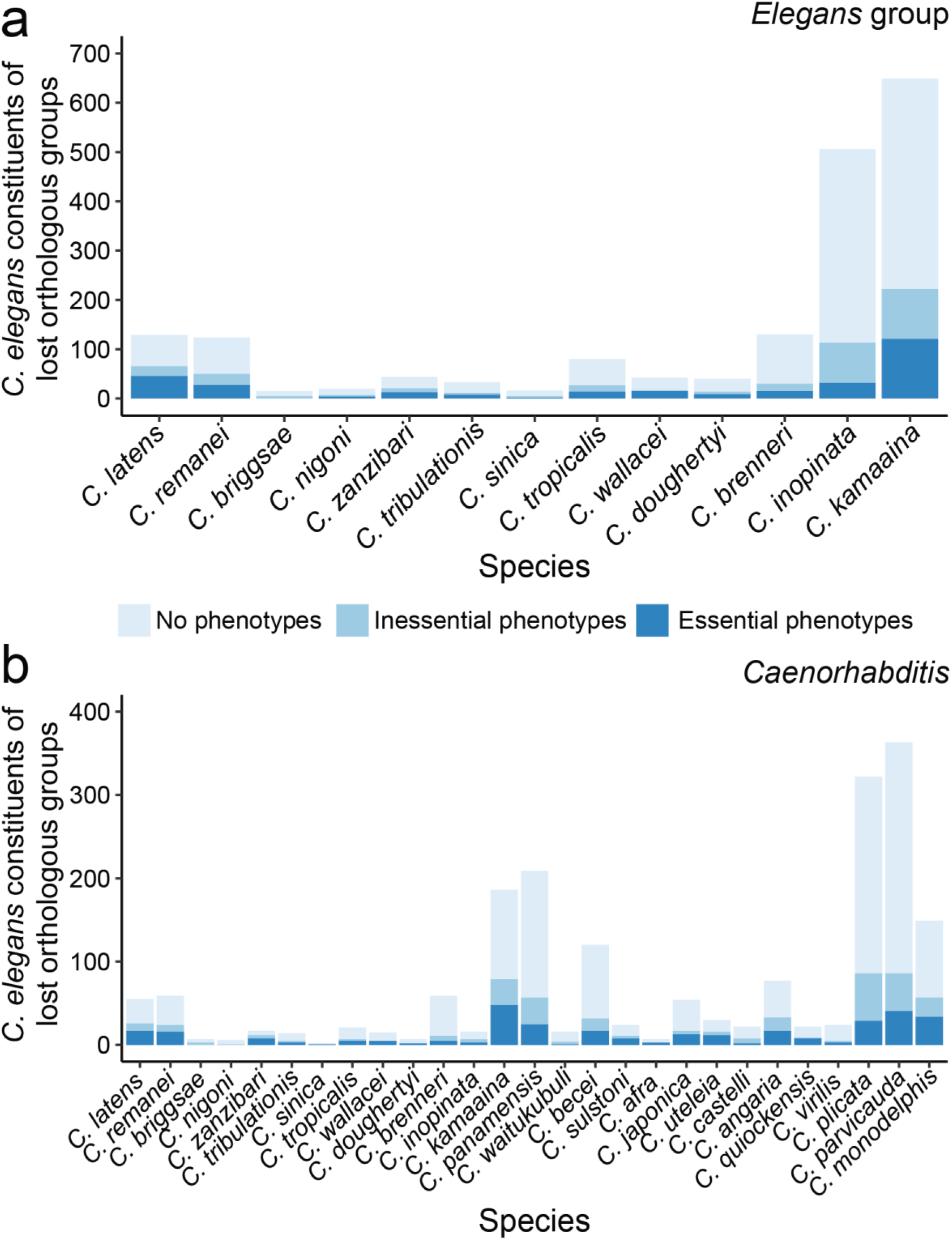
Patterns of ortholog loss reveal rampant developmental system drift in *Caenorhabditis*. Plotted are the number of genes with essential, inessential, or no WormBase phenotype terms among the *C. elegans* gene constituents of lost orthologous groups when considering the *Elegans* group (a) or all *Caenorhabditis* (b). Species are roughly ordered by phylogenetic relatedness. *C. elegans* is not plotted because orthologous groups lost only in *C. elegans* have no WormBase annotations.

### Patterns of pleiotropy, specificity, and spatiotemporal transcript abundance among lost genes

Do lost genes share any common features, or can gene loss be predicted? To address this question, other features of *C. elegans* genes associated with species-specific lost orthologous groups were also examined. In addition to phenotypes, WormBase also contains “anatomy” and “life stage” ontologies. These relate spatial (“anatomy”) and temporal (“life stage”) expression patterns to genes. WormBase also contains information about pairwise interactions among genes (“interaction”), which are defined by epistatic genetic interactions or physical/biochemical interactions. WormBase also tracks the number of peer-reviewed scientific papers that mention a given gene as a “reference count.” Additionally, the domains in all *C. elegans* proteins were defined using the 16,713 Pfam domain seed alignments and HMMER (Finn, et al. 2015). The per-gene number of unique features of all of these categories (phenotype, tissue, life stage, interaction, reference count, and domain) were counted. This then provides quantitative measures of: the consequential phenotypic complexity upon perturbation of a given gene (phenotype); the extent of expression specificity across space and time of a given gene (anatomy and life stage); the connectedness of a given gene in its biological network (interaction); the extent to which a given gene has been studied (reference count); and the number of functional modules a given gene has (domain). Taken together, these provide various coarse measures of genetic specificity and pleiotropy.

All *C. elegans* genes associated with species-specific lost orthologous groups, irrespective of species, were denoted as lost at the two levels of phylogenetic scope (*Elegans* group or all *Caenorhabditis*). In addition, genes lost only in *C. inopinata* (within the context of the *Elegans* group; Figure 2a and Figure 3a) were also addressed. *C. inopinata* is a species worthy of consideration on its own for two major reasons. First, it is the closest known relative of *C. elegans* (Kanzaki, et al. 2018; Woodruff, et al. 2018) and represents the lower bound of phylogenetic distance from the reference species among the organisms in this study. Second, it is morphologically and ecologically divergent from *C. elegans*, and genes lost only in *C. inopinata* may be good candidates for understanding the genetic basis of morphological and ecological divergence (Kanzaki, et al. 2018; Woodruff and Phillips 2018; Woodruff, et al. 2018). In any case, at all levels of phylogenetic consideration, genes that were not denoted as “lost” were called “retained.” Then, the distributions of numbers of WormBase phenotypes, anatomies, life stages, interactions, reference counts, and PFAM domains among lost and retained genes at the three levels of phylogenetic scope were compared.

The total number of *C. elegans* genes from lost orthologous groups is substantial when considering both the *Elegans* group (1,828 or 9.0% of *C. elegans* protein-coding genes) and all *Caenorhabditis* (1,903 or 9.4%). Furthermore, although these genes represent similar proportions of the genome, they largely do not overlap (only 464 genes are shared among the two groups (464/3,267 or 14.2%); Supplemental Figure 16). In the case of only *C. inopinata*, the number of lost *C. elegans* genes is far less (506 or 2.5%). The distribution of ontological term numbers among lost and retained genes described above also varies depending on the phylogenetic scope (Figure 4a). Among genes lost only in *C. inopinata*, the average number of all WormBase terms and domains are significantly lower than among those genes that are retained (Figure 4a). When considering genes lost among *Elegans* group members, this pattern is similar, although the effect size of gene loss is far less across all categories (Figure 4a). However, when looking at the broadest phylogenetic level, all *Caenorhabditis*, this pattern is largely eroded. Lost genes at this level are largely no different from retained genes; however, lost genes reveal a subtle but detectable increase in the number of domains relative to retained genes (Cohen’s *d* effect size=0.073; 95% confidence interval=0.019-0.13; Mann-Whitney U *p*=2.09 × 10^−10^, W=11584369). Thus, the degree of specificity and pleiotropy among lost genes depends upon the phylogenetic context considered. At narrower phylogenetic scopes, lost genes tend to be less pleiotropic and specific than retained genes, and as the phylogenetic scope broadens, lost genes tend to resemble retained genes.

**Figure 4.**
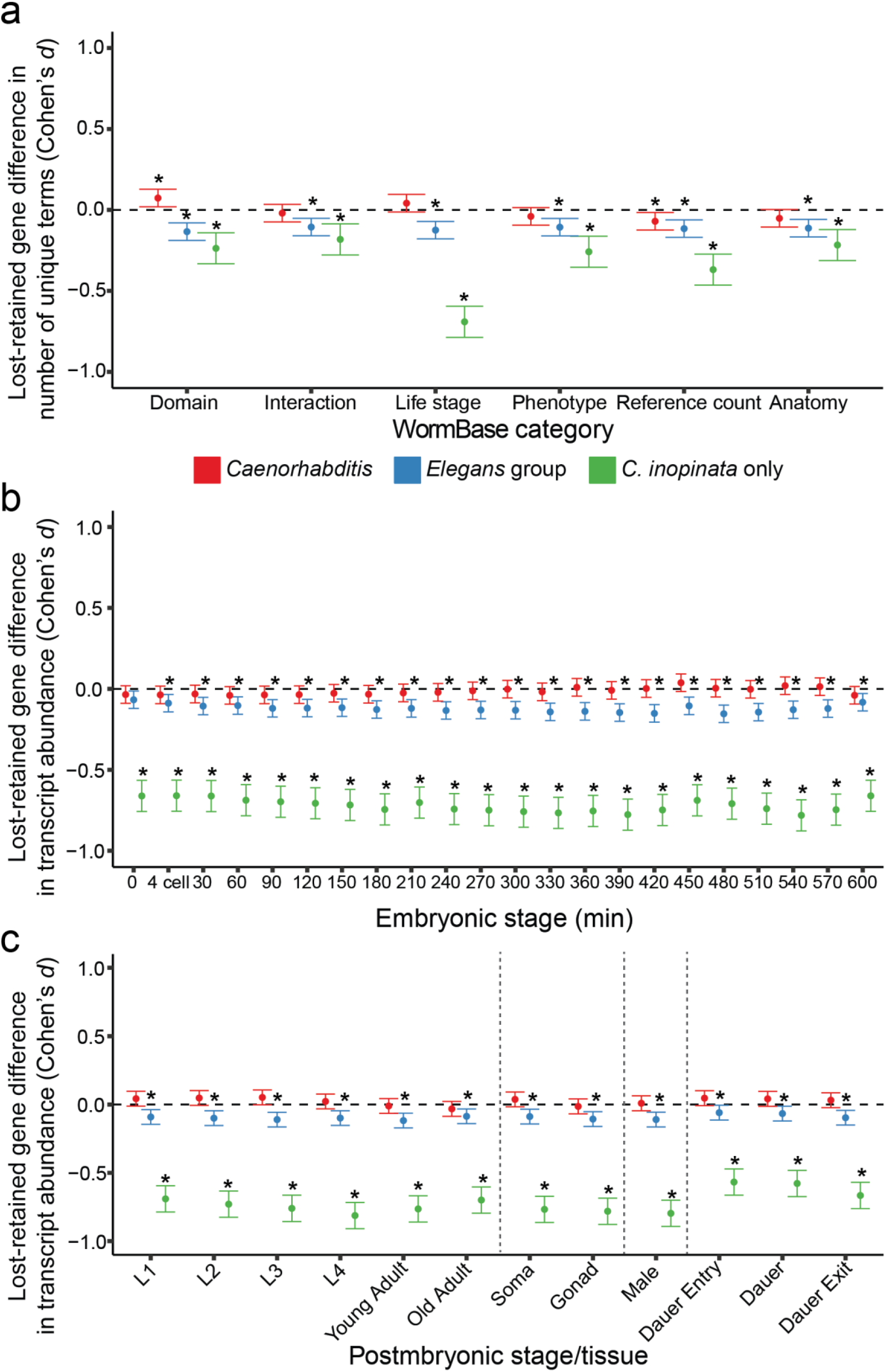
The impact of pleiotropy and transcription on gene loss depends on phylogenetic scope. In all panels the effect size (Cohen’s *d*) of gene loss relative to gene retention is plotted. Here, all species-specific gene losses are pooled and denoted as lost genes; *C. elegans* measures among lost and not lost (i.e., retained) genes are being compared. An effect size of −1 notes that the average value among lost genes is one pooled standard deviation lower than retained genes; an effect size of 0 (dashed horizontal line) reveals on average no difference between lost and retained genes. Error bars represent 95% confidence intervals of the effect size. Distributions of all categories for lost and retained genes across all levels of phylogenetic scope can be found in Supplemental Figures 17-25. In all panels, * = Mann-Whitney U p < 0.01 in a comparison of lost and retrained genes. (a) The effect size of gene loss on WormBase feature number per gene. Numbers of domains were determined with HMMER. All other features were retrieved from WormBase. (b) The effect size of gene loss on transcript abundance (1+log_2_(dcpm)) per gene across stages of embryonic development. All stages were recorded as minutes past fertilization with the exception of “4 cell;” the four cell stage is typically ~30 minutes past fertilization in typical rearing conditions (Altun, et al. 2002-2016). (c) The effect size of gene loss on mean transcript abundance (1+log_2_(dcpm)) per gene across stages of postembryonic development. Vertical dotted lines separate hermaphroditic postembryonic stages, hermaphroditic adult soma and germ line preparations, male, and dauer-related stages. RNAseq data were retrieved from (Boeck, et al. 2016).

In addition to the ontological information accessible in WormBase, the transcriptome of *C. elegans* has also been intensively studied (Gerstein, et al. 2010; Levin, et al. 2012; Hashimshony, et al. 2015). One recent report measured transcript levels at 30-minute intervals across embryonic development to define the “time-resolved transcriptome of *C. elegans*” (Boeck, et al. 2016). In addition to patterns of transcription across embryogenesis, this study also included measures of gene expression across various postembryonic stages (Boeck, et al. 2016). To provide further biological context to the lost genes defined above, the transcriptomic data from the Boeck et al. study was paired with this information (Figures 4b-c). Much like with the WormBase ontological terms (Figure 4), the transcriptional abundances of lost and retained genes at various embryonic (Figure 4b) and postembryonic (Figure 4c) stages were compared within the context of three phylogenetic levels (only *C. inopinata*, the *Elegans* group, and all *Caenorhabditis* (Figure 4b-c)). Like the WormBase ontological terms, patterns of gene expression among lost genes varied depending on the phylogenetic scope. Among genes lost only in *C. inopinata*, lost genes exhibited much lower expression than retained genes across all developmental stages (average effect size= −0.72; Figure 4b-c). Among genes lost in *Elegans* group species, lost genes are only slightly less expressed than retained genes at all developmental stages (average effect size= −0.11; Figure 4b-c). And, as with the WormBase terms, no differences in expression among lost and retained genes could be detected at any developmental stage at the broadest phylogenetic scope (all *Caenorhabditis*; Figure 4b-c). Thus, transcriptional abundance among genes with the capacity to be lost also depends on phylogenetic context, and at narrower phylogenetic scopes, lost genes tend to have lower expression than retained genes.

Multivariate analyses were also performed to test the impact of transcriptional activity and the number of WormBase ontology terms on gene loss. All models of gene loss are significant when including all transcription and ontology count variables simultaneously (MANOVA: all *Caenorhabditis* p= 5.5 × 10^−7^, Pillai’s trace=0.0050; *Elegans* group p=2.8 × 10^−13^, Pillai’s trace=0.0072; *C. inopinata* only p<2.2 × 10^−16^, Pillai’s trace=0.021; see supplemental data). The most important contributors to gene loss (i.e., the factors with the largest coefficients) depend on the phylogenetic scope. For all *Caenorhabditis*, these are transcription at 330 minutes-past-fertilization (mpf; Linear Discriminant 1 (LD1) coefficient= −0.49), adult soma-specific expression (LD1 coefficient= 0.44), and transcription at 300 mpf (LD1 coefficient=0.38; see supplemental data). For the *Elegans* group, these are young adult hermaphrodite expression (LD1 coefficient=-0.38), transcription at 420 mpf (LD1 coefficient= −0.36), and transcription at 300 mpf (LD1 coefficient=0.35; see supplemental data). And for *C. inopinata* only, these are transcription at 480 mpf (LD1 coefficient=0.24), L2 expression (LD1 coefficient=0.22), and transcription at 390 mpf (LD1 coefficient= −0.22; see supplemental data). However, the proportion of the variance explained of these models is small (all *Caenorhabditis*, logistic regression pseudo-*R*^2^=0.013; *Elegans* group, logistic regression pseudo-*R*^2^=0.019), although the models perform better for genes lost only in *C. inopinata* (logistic regression pseudo-*R*^2^=0.12). These are consistent with the high overlap of lost and retained genes in principal components (Supplemental Figures 26-28) and linear discriminant (Supplemental Figure 29) space. Thus there is a generally weak but detectable impact of transcriptional activity and gene ontology count on gene loss that increases as the phylogenetic scope narrows.

## Discussion

### Widespread turnover of developmental genetic systems in a group with highly conserved morphology

Gene loss is a widespread driver of phenotypic change (Albalat and Cañestro 2016). At the same time, genetic perturbations such as gene loss often have no discernable phenotypic effects, underscoring the roles of genetic redundancy and context-dependence in phenotype generation. Here, I explored the extent of potential gene loss in *Caenorhabditis* nematodes by describing orthologous genes that are present (or detectable) in all species but one at a given phylogenetic level. I then situated these genes within their functional context by connecting them to known phenotypic roles through *C. elegans* WormBase ontologies and transcriptional data (Boeck, et al. 2016). What can these patterns reveal about the evolution of developmental systems?

*Caenorhabditis* species are notable for their morphological constancy in the face of tremendous genetic divergence (Kiontke, et al. 2004) (although the fig-associated *C. inopinata* is morphologically exceptional (Kanzaki, et al. 2018; Woodruff, et al. 2018)). Within the *Elegans* group, species are largely morphologically indistinguishable (although there are male tail features that distinguish some clades), and mating tests or molecular barcoding is usually necessary to delineate groups (Kiontke, et al. 2011; Félix, et al. 2014; Stevens, et al. 2019). This morphological conservation also holds throughout development—the pattern of cell divisions from the single-cell zygote to the reproductive adult is also largely invariant among species (Zhao, et al. 2008; Memar, et al. 2018). This conserved developmental pattern persists despite high genetic divergence; *Caenorhabditis* species span a genetic distance comparable to that between mice and lampreys (Kiontke, et al. 2004). Here, we document widespread species-specific gene loss, with dozens of genes being lost on average per species (Fig. 2–3). These patterns are consistent with previous observations of sequence-level (Cutter 2008) and gene copy number (Stevens, et al. 2019) variation among *Caenorhabditis* species, with rampant genetic turnover underlying a stable developmental and morphological system.

Notably, only 36% of gene losses observed in other species are associated with obvious phenotypes in *C. elegans*, with up to 20% of these (on average) actually being essential for viability and fecundity. How can genes presumably essential for a conserved developmental system be lost so often? The phenotype terms used to functionally annotate these orthologs are derived from the vast background knowledge of *C. elegans* model systems genetics and are generally informed by mutation or RNAi evidence (Schindelman, et al. 2011). So one possible explanation for this pattern could be that the developmental genetics of *C. elegans* are exceptionally idiosyncratic such that the functional annotations derived from this species are not be widely applicable across the genus. In this case, interpretations regarding the loss of essential genes would be mistaken because *C. elegans* might just be an unusual species; that is, it is possible that in most *Caenorhabditis* lineages these orthologous groups are indeed dispensable, and their essential functions are novel to or derived in *C. elegans*. Several lines of evidence lend some credibility to this explanation. For one, *C. elegans* is a self-fertile hermaphrodite, a mode of reproduction that is largely the exception in this group, as only three species in the genus have hermaphrodites (most are male/female) (Stevens, et al. 2019). These species represent 11% (3/28) of the assemblies used in this study. As the evolution of selfing has profound consequences for multiple aspects of an organism’s biology (Thomas, et al. 2012), this could promote an idiosyncratic developmental system not comparable to its close relatives. Further, most of the evidence used for these phenotype terms are derived from studies using the N2 strain of *C. elegans*, which is thought to be a laboratory domesticated strain (Sterken, et al. 2015). As *C. elegans* N2 is biologically exceptional with respect to *C. elegans* as a species (Sterken, et al. 2015), it may not be representative of the genus as a whole. *C. elegans* is also divergent in its regulation of small RNAs—there is ample variation in susceptibility to RNAi by feeding in *Caenorhabditis* and *C. elegans* is particularly vulnerable (Nuez and Félix 2012). Thus *C. elegans* may represent an idiosyncratic developmental system whose annotations belie a misleading interpretation of functional gene loss. Future studies utilizing whole-genome approaches to genetic function in other *Caenorhabditis* species (Verster, et al. 2014) will help to inform the extent of functional diversity among orthologous genes in this group.

Nevertheless, while it is formally possible that *C. elegans* has a particularly idiosyncratic biology, a much more plausible explanation for these patterns of gene loss is rampant developmental system drift. Developmental systems drift occurs when divergent developmental programs underlie otherwise conserved morphological features (True and Haag 2001). One reason to suspect *C. elegans* is comparable to is close relatives is that many orthologs do maintain conserved function across the genus (Haag, et al. 2018), and some genes have deeply conserved functions through nematode phylogeny (Crook 2014; Haag, et al. 2018; Kasimatis and Phillips 2018). Thus at least some aspects of the *Caenorhabditis* genetic system are conserved and underlie a static morphology. Instead, the combination of many genes being lost in a species-specific manner while being essential for fitness in at least one species is consistent with widespread, species-specific turnover of genetic function despite morphological stasis.

This evolutionary pattern of heterogeneity in essential gene function is consistent with other more direct functional assays across species. *C. elegans* was among the first metazoans to be interrogated with genome-wide genetic knockdown via RNAi (Fraser, et al. 2000; Gönczy, et al. 2000; Piano, et al. 2000; Maeda, et al. 2001; Kamath, et al. 2003). Since then, dozens of such screens have been implemented (E Yanos, et al. 2012), which provides a comparatively exhaustive picture of genetic function within this species. The application of a similar approach in a close relative, the hermaphroditic *C. briggsae*, affords an opportunity to test the extent of functional conservation across orthologous genes directly (Verster, et al. 2014). In this case, over 25% of orthologous genes have divergent functions between the two species, consistent with widespread functional turnover and developmental system drift (Verster, et al. 2014). This point is further emphasized by a genome-wide RNAi screen among *C. elegans* wild isolates by (Paaby, et al. 2015), who found widespread variation in maternal-effect gene knockdown penetrance suggestive of a developmental system in flux within as well as between species. This is perhaps best exemplified in the development of the nematode vulva, which has long been the study of detailed genetic analysis and which displays highly divergent developmental processes and genetic pathways despite yielding a highly conserved morphological structure across nematode phylogeny (Haag, et al. 2018). The picture of developmental systems drift that is emerging within nematodes is consistent with observations from other studies, such as variation in postzygotic isolating factors among closely-related species (including fruit flies, mammals, birds, butterflies, monkeyflowers, and other taxa (Coyne and Orr 2004)) and vertebrate limb development (Shubin and Alberch 1986; Haag and True 2018). Overall, the patterns of gene loss observed here are consistent with a body of evidence detailing a variety of surprisingly dynamic developmental processes across the tree of life.

### Predicting gene loss with transcription and pleiotropy

Functional phenotypic annotations revealed that lost genes often have essential functions in *C. elegans*. Can additional information about these genes be used to predict gene loss? I retrieved WormBase ontology terms (Lee, et al. 2017), Pfam domains (Finn, et al. 2015), and stage-specific transcriptional abundance data (Boeck, et al. 2016) to examine if they can differentiate gene retention from gene loss. With respect to WormBase ontology terms, the number of unique terms per feature for each gene was used and provides crude metrics for the extent of its pleiotropy (i.e., the number of phenotypes a gene has or the number of tissues and/or developmental stages a gene is expressed in).

Intuitively, one might expect less widely expressed and less pleiotropic genes to be less constrained by selection and more prone to loss. From a univariate perspective, the impact of transcriptional abundance and pleiotropy on gene loss varies by phylogenetic scope (Figure 4). Patterns in genes lost only in *C. inopinata* and in the *Elegans* group largely agreed with these expectations—retained genes were more likely to be expressed across all developmental stages (Fig. 4b-4c) and have more WormBase ontology features than lost genes. However, this did not hold for genes lost when considering the genus as a whole. Surprisingly, genes lost at these different levels of phylogenetic consideration largely did not overlap (Supplemental Figure 16) and revealed different patterns of differential transcriptional abundance. Specifically, genes lost with respect to the *Elegans* group had significant but small effects on transcriptional abundance across development when compared to retained genes (Figure 4b-c). Conversely, genes lost with respect to *Caenorhabditis* were not distinguishable from retained genes in transcriptional patterns (Figure 4b-c). These patterns are largely mirrored in the WormBase term metrics (Figure 4a). As genes lost only in *C. inopinata* exhibited moderately low transcription across the board (Fig. 4b-c), this suggests that as the phylogenetic scope broadens, the impact of transcription and pleiotropy (in a single reference species) on gene loss weakens. This is also consistent with widespread developmental system drift and the rapid evolution of developmental processes.

When present, these differences in transcriptional abundance appear to span broad periods of developmental time, and gene loss at broader phylogenetic levels has miniscule or no effects on these traits (Fig. 4b-c). In a principal component analysis, retained and loss genes do not overlap in multidimensional space at any level of phylogenetic scope (Supplemental Figures 26-28). Furthermore, linear discriminant analysis, whose aim is to find the function that best separates two groups, is also unable to distinguish lost and retain genes (although prediction is marginally better in the case of genes lost only in *C. inopinata* (Supplemental Figure 29)). Thus, despite subtle, broad detectable differences in transcriptional abundance at discrete time points (Fig. 4b-c; supplemental data), these data cannot predictably distinguish genes with a tendency to be lost in a species-specific manner, consistent with pervasive turnover of developmental mechanisms along nematode phylogeny. And although gene loss is difficult to predict in this context, it is possible that additional biological information (such as those uncovered in the modENCODE project (Gerstein, et al. 2010)) could be harnessed to understand how and why genes are lost in this manner.

### Caveats

Here, I have set to define and understand patterns of gene loss across *Caenorhabditis* nematode species with publicly available genome assemblies, and gene losses were inferred through a common computational pipeline applied to these assemblies and their associated protein sets. A potential complicating factor in the interpretation of these results is variation in genome assembly and annotation quality. There is clearly variation in both assembly and annotation quality in the genomes used in this study (Supplemental Figures 1-5). All genomes in this study used RNAseq data to inform their annotations (Howe, et al. 2016; Kanzaki, et al. 2018; Yin, et al. 2018; Stevens, et al. 2019) and provide evidence-based approaches to bolster the reliability of their protein sets. Despite this, questions remain regarding annotation quality. For instance, *C. sinica* has a notably large protein set with 34,696 coding genes. Inflated gene copy number due to collapsed alleles is a known problem with such hyperdiverse gonochoristic species (Barrière, et al. 2009), and it is possible that this reflects an overestimate of gene number in this case. However, overestimates of gene number should not impact inferences of gene loss per se, as there is no reason to think collapsed alleles would cause biases against annotating genes that are present. Additionally, BUSCO completeness scores, which are measured by comparing protein sets against a set of proteins thought to be largely universal among certain organismal groups (Simão, et al. 2015), reveal the *C. angaria* protein set as an outlier with a 63.5% completeness score. This is suggestive of an incomplete protein set which would cause overestimates of gene loss in this species. Thus, particularly for genomics with low completeness metrics, these are likely to be overestimates of the extent of gene loss in this group.

Variation in genome assembly quality is more problematic for this study. Only four of the 28 species used in this study have chromosome level assemblies, and most of the assemblies are highly fragmented (Supplemental Figures 3-4). Although our computational pipeline can presumably overcome shortcomings in annotations through genomic alignments, it cannot account for genomic regions that have not been assembled. Thus a major caveat of this work is that these specific inferences of gene loss are provisional due to the high variation in completeness among the genome assemblies used here. Future work using chromosome-level assemblies will be required to ascertain more precise estimates of gene loss in this group. That said, these estimates are not without value—most of the assemblies used here have high completeness metrics (Supplemental Figure 5) and chromosome-level completeness would likely not have much impact on the qualitative interpretation that essential genes are often lost. This is consistent with essential genes being lost even in the assemblies with chromosome-level completeness (Figure 3). So although the quantitative extent of gene loss per species may be overestimated, the pervasiveness of developmental system drift remains a reasonable interpretation.

Additionally, the method of orthology assignment itself may impose biases upon inferences of species-specific loss. Here, loss was defined as being present in every species but one, given some phylogenetic scope. If there is rapid clade-specific genetic divergence, distance-based clustering may lead to the splitting of orthologous groups. There are thousands of orthologous groups that are restricted to a few species (see Supplemental Figure 6 for the example of orthologous groups found only in *C. remanei* and *C. latens*) or are species-specific. Presumably this can be partly explained by the emergence of clade-specific genes, but this could also be due to rapid clade-specific divergence. These types of orthologs would be excluded from this analysis and could actually underestimate the extent of gene loss. Additionally, as the method of orthologous group inference begins with predicted protein sets, genes that are erroneously unidentified in multiple species would not be included here, also underestimating the amount of gene loss. And finally, there is the implicit use of parsimony in assuming gene loss throughout this study. In all cases gene loss is assumed because orthologs are present in all other species; there remains the possibility of multiple gene gains for any of these orthologs, although this parsimony issue is likely not affecting interpretations.

## Conclusions

*C. elegans* is a widely used model system for biomedical genetics, and it has been at the forefront of metazoan genomics since the inception of the discipline. However, the organisms and resources needed to place these findings in their broader evolutionary and phylogenetic contexts are only recently becoming available. Here, the previously-sequenced genomes of 28 *Caenorhabditis* species revealed that all have lost genes that perform essential functions in *C. elegans*, suggesting that developmental processes are rapidly evolving in this group. As presumably essential genes are turning over rapidly, this also suggests that biological functions among conserved genes may also be changing quickly. This underscores the need of comparative approaches in interpreting and translating findings in model systems genetics across large evolutionary distances.

## Materials and Methods

### Determining species-specific gene loss

28 *Caenorhabditis* protein sets and genome assemblies were retrieved from the *Caenorhabditis* Genomes Project (Slos, et al. 2017; Stevens, et al. 2019)(caenorhabditis.org; *C. afra*, *C. castelli*, *C. doughertyi*, *C. inopinata* (formerly *C.* sp. 34), *C. kamaaina*, *C. latens*, *C. monodelphis*, *C. plicata*, *C. parvicauda* (formerly *C.* sp. 21), *C. zanzibari* (formerly *C.* sp. 26), *C. panamensis* (formerly *C.* sp. 28), *C. becei* (formerly *C.* sp. 29), *C. utelei* (formerly *C*. sp. 31), *C. sulstoni* (formerly *C*. sp. 32), *C. quickensis* (formerly *C*. sp. 38), *C. waitukubuli* (formerly *C*. sp. 39), *C. tribulationis* (formerly *C*. sp. 40), *C. virilis*) and WormBase Parasite (Howe, et al. 2016)(parasite.wormbase.org; *C. angaria*, *C. brenneri*, *C. briggsae*, *C. elegans*, *C. japonica*, *C. remanei*, *C. sinica*, *C*. *tropicalis*), or were otherwise shared (*C. nigoni* (Yin, et al. 2018) and *C. wallacei*, E. Schwarz pers. comm.). The *Diploscapter coronatus* genome (Hiraki, et al. 2017) was used as an outgroup. See Supplemental Figures 1-5 for measures of assembly size, gene number, scaffold number, N50, and completeness of these retrieved genome projects.

Alternative splice variants were removed from the protein sets such that each protein-coding gene was represented by the longest-isoform protein. To identify orthologous groups, 841 all v. all blastp searches (Camacho, et al. 2009) were performed among the protein sets (version BLAST+ 2.6.0; blastp options -outfmt 6 -evalue 0.001 -num_threads 8). The blastp results were then fed into OrthoFinder (Emms and Kelly 2015)(version 1.1.8; options -b -a 10) to define orthologous proteins. To determine species-specific gene losses, orthologous groups that were present in every species but one were extracted. It is important to emphasize that throughout this paper, “loss” will be assumed to be equivalent to this type of species-specific absence, as opposed to many other possible patterns of repeated loss in multiple species; here we are only examining patterns of species-specific loss. Additionally, as these orthologous groups were absent only in one species, loss is the most parsimonious explanation for their absence as opposed to multiple independent gains in the other species. This analysis was performed at two phylogenetic levels (all *Caenorhabditis* species and only *Elegans* group species (Figure 1)) as the number of shared orthologous groups decreases with phylogenetic distance (Supplemental Figure 6).

To be conservative in estimating the extent of species-specific gene loss, additional filters were applied to the OrthoFinder results. OrthoFinder implements a size-normalization step to BLAST bit scores in order to account for the correlation between protein length and bit score (Emms and Kelly 2015). Concerned that poor gene models that inflate gene length would distort the proper inference of orthologous groups, species-specific orthologous group losses as defined by OrthoFinder were re-examined for best-reciprocal blastp hits with *C. elegans* among the results described above. If putative losses were revealed to have a best reciprocal blastp hit with *C. elegans*, they were removed from consideration as such a species-specific gene loss. Furthermore, as OrthoFinder uses predicted proteins to define orthologous groups, unannotated genes may be spuriously determined as species-specific losses. To address this problem, the *C. elegans* protein constituents of putative losses were aligned to their respective genome assemblies using tblastn (Camacho, et al. 2009)(version BLAST+ 2.5.1; options -outfmt 6), which searches entire translated genomes without the need of gene models. If a putative lost *C. elegans* protein had a best tblastn hit outside of a predicted coding gene in the respective genome assembly (using the BEDtools (Quinlan 2014) *intersect* function (version 2.25.0; option -v) with the respective genome assembly’s annotations to retrieve alignments that fall outside of predicted protein-coding regions), this ortholog was then removed from consideration as being a species-specific loss. This pipeline then accounts for problems incurred by poor gene annotations which OrthoFinder cannot address.

### Connecting species-specific ortholog losses to WormBase ontology, Pfam domain, and RNAseq data

To understand the functional and biological characteristics of lost genes, the *C. elegans* members of genes lost in all non-*C. elegans* species (at both levels of phylogenetic consideration; Figure 1) were extracted. These were then paired with WormBase (Howe, et al. 2015) ontology information (specifically phenotype, anatomy, life stage, and reference count), which were retrieved with the SimpleMine tool (Howe, et al. 2015) with all *C. elegans* protein-coding genes. WormBase is a database housing information regarding the genetics, genomics, and general biology of *C. elegans* and other nematodes. Particularly, it has collected from the literature and scientific community genome-wide, gene-specific, and hand-curated information including: the biological consequences of mutation and RNAi exposure (“phenotype”); tissue-specific expression patterns (“anatomy”); temporal expression patterns (“life stage”); genetic and biochemical interactions with other genes and their encoded proteins (“interaction”); and its number of scientific papers (“reference count”), among other features (Howe, et al. 2015). These features have been formalized as genomic ontologies (Lee and Sternberg 2003; Schindelman, et al. 2011) and were used in this study. Essential genes were defined as any of those with WormBase phenotypes containing the words “lethal,” “arrest,” “sterile,” or “dead.” This included 58 unique phenotype terms (see supplemental data for the list of essential phenotypes).

Domains were identified in the *C. elegans* protein set using all domains defined by the Pfam database (Finn, et al. 2015). Seed alignments for the domains were retrieved from the Pfam FTP site (ftp://ftp.ebi.ac.uk/pub.databases/Pfam/current_release/Pfam-A.seed.gz), and hidden markov models were constructed with HMMER (version 3.1b2; function *hmmbuild*) (Eddy 1998) using default parameters. Then, the models were used to search for all domains across all *C. elegans* proteins using HMMER (function *hmmsearch* option –tblout) using default parameters. These results were then used to determine the number of domains per protein-coding gene in *C. elegans*.

In addition, stage-specific quantitative RNAseq data was also paired with the *C. elegans* members of genes lost in all non-*C. elegans* species. Data from Boeck et al. 2016 (Boeck, et al. 2016), which examined transcript abundance at 30 min intervals of embryonic development in *C. elegans*, was used to capture expression dynamics and the potential for predicting gene loss. This data set also included expression data for postembryonic larval stages, dauer developmental stages, males, the soma, and the germ line. Here, the mean depth of coverage per million across biological replicates was taken as the representative transcript level for a gene at a given stage. And as only 19,712 protein-coding genes were reported as being expressed in this data set, only these genes were included in subsequent analyses here.

All statistics were performed using the R programming language. Mann-Whitney U tests, principal components analyses, and MANOVA tests were performed with the base functions *wilcoxon.test*, *prcomp*, and *manova*. Cohen’s *d* effect sizes were estimated using the *cohen.d* function in the “effsize” package (https://github.com/mtorchiano/effsize/). Linear discriminant analyses were performed using the *lda* function in the “MASS” package (Venables and Ripley 2013), and discriminant functions were projected back onto the data using the *predict* function. Multiple logistic regression models were performed with the *lrm* function in the “rms” package http://biostat.mc.vanderbilt.edu/wiki/Main/Rrms).

## Supporting information

Elegans group lost genes

Genes lost only in C. inopinata

WormBase and transcriptomic data

Statistics summaries

List of essential phenotypes

Caenorhabditis lost genes

## Supplementary Material

Supplemental Figure 1. Assembly sizes.

Supplemental Figure 2. Number of protein coding genes.

Supplemental Figure 3. Scaffold numbers.

Supplemental Figure 4. N50 values.

Supplemental Figure 5. BUSCO completeness scores.

Supplemental Figure 6. The number of recovered shared orthologous groups decreases as more species are included.

Supplemental Figure 7. The distribution of species-specific lost orthologous groups among all *Caenorhabditis* and only the *Elegans* group.

Supplemental Figure 8. BUSCO completeness score and the number of lost orthologous groups when considering the *Elegans* group.

Supplemental Figure 9. BUSCO completeness score and the number of lost orthologous groups when considering all *Caenorhabditis*.

Supplemental Figure 10. Scaffold number and the number of lost orthologous groups when considering the *Elegans* group.

Supplemental Figure 11. Scaffold number and the number of lost orthologous groups when considering all *Caenorhabditis*.

Supplemental Figure 12. N50 and the number of lost orthologous groups when considering the *Elegans* group.

Supplemental Figure 13. N50 and the number of lost orthologous groups when considering the all *Caenorhabditis*.

Supplemental Figure 14. The average gene copy number per orthologous group is low.

Supplemental Figure 15. The distribution of orthologous group gene copy numbers less than five.

Supplemental Figure 16. The *C. elegans* gene constituents of species-specific lost orthologous groups do not largely overlap among different levels of phylogenetic consideration.

Supplemental Figure 17. The distribution of number of unique WormBase terms or Pfam domains among retained and lost genes when considering all *Caenorhabditis*.

Supplemental Figure 18. The distribution of number of unique WormBase terms or Pfam domains among retained and lost genes when considering the *Elegans* group.

Supplemental Figure 19. The distribution of number of unique WormBase terms or Pfam domains among retained and lost genes when considering only *C. inopinata*.

Supplemental Figure 20. The distribution of transcriptional abundance by embryonic stage among retained and lost genes when considering all *Caenorhabditis*.

Supplemental Figure 21. The distribution of transcriptional abundance by embryonic stage among retained and lost genes when considering the *Elegans* group.

Supplemental Figure 22. The distribution of transcriptional abundance by embryonic stage among retained and lost genes when considering only *C. inopinata*.

Supplemental Figure 23. The distribution of transcriptional abundance by postembryonic stage among retained and lost genes when considering all *Caenorhabditis*.

Supplemental Figure 24. The distribution of transcriptional abundance by postembryonic stage among retained and lost genes when considering the *Elegans* group.

Supplemental Figure 25. The distribution of transcriptional abundance by postembryonic stage among retained and lost genes when considering only *C. inopinata*.

Supplemental Figure 26. Principal components analysis, all *Caenorhabditis*.

Supplemental Figure 27. Principal components analysis, *Elegans* group.

Supplemental Figure 28. Principal components analysis, only *C. inopinata*.

Supplemental Figure 29. Linear discriminant analysis of transcriptomic and WormBase data.

Supplemental Table 1. The top 25 most common phenotypes in WormBase.

Supplemental Table 2. The top ten “most pleiotropic” genes as measured by number of unique WormBase phenotypes.

Supplemental Table 3. The top ten most widely studied genes as measured by WormBase reference count.

Supplementary data (essential_phenotypes.txt; lost_gene_list_all_caenorhabditis.tsv; lost_gene_list_elegans_group.tsv; lost_gene_list_inopinata.txt; lost_genes_wormbase_boeck.tsv; statistics_summaries.xlsx;) are available at *Journal*.

## Data deposition and accessibility

Genome sequences, protein sets, and annotations were retrieved from the *Caenorhabditis* Genomes Project (caenorhabditis.org) or WormBase ParaSite (parasite.wormbase.org). Data files and code associated with this study have been deposited in Github at https://github.com/gcwoodruff/gene_loss.

## Acknowledgements

I thank Mark Blaxter, Erich Schwarz, Janna Fierst, Janet Young, Matthew Rockman, and Patrick Phillips for sharing genomic data. Bill Cresko and his laboratory members provided helpful comments throughout the development of this work. Patrick Phillips and Erich Schwarz provided valuable feedback on earlier versions of this manuscript. This work was supported by funding from the National Institutes of health to GCW (5F32GM115209-03) and to Patrick Phillips (R01 GM-102511; R01 AG049396).

