## Supplemental Figures for "Patterns of putative gene loss suggest rampant developmental system drift in nematodes"

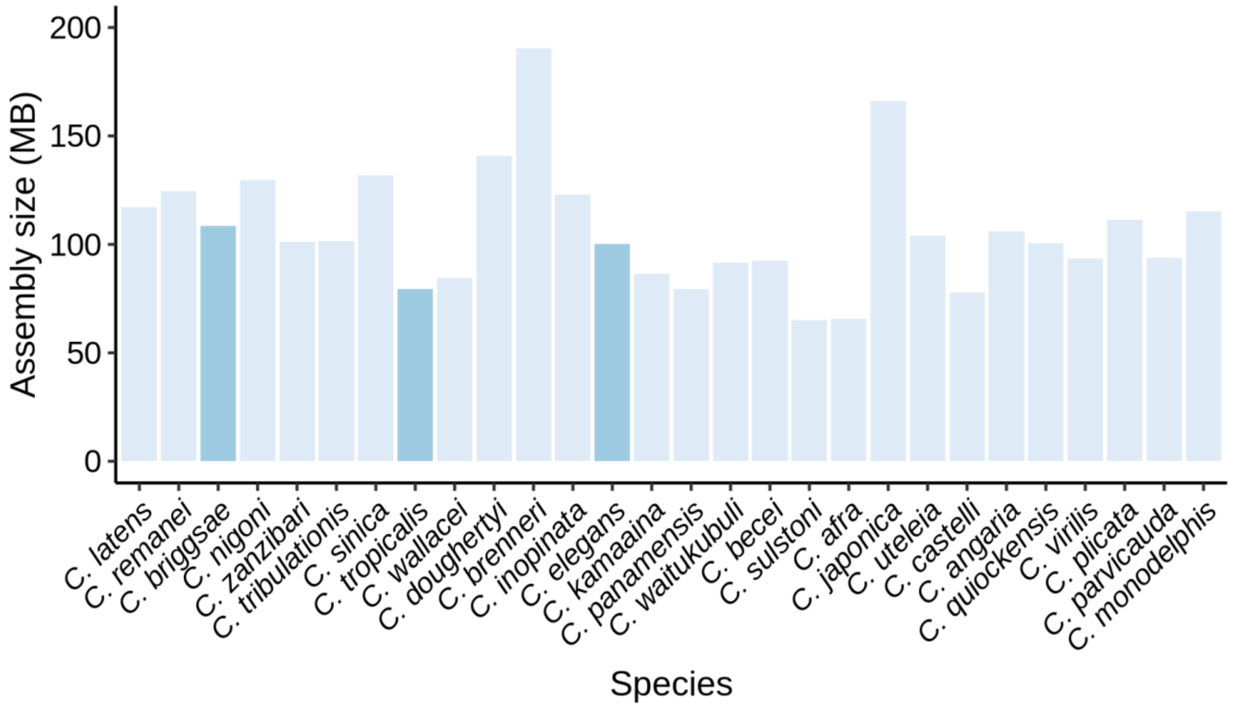


Supplemental Figure 1. Assembly sizes of the genomes used in this study. Species are highlighted by reproductive mode (light blue, female/male; dark blue, hermaphroditic).

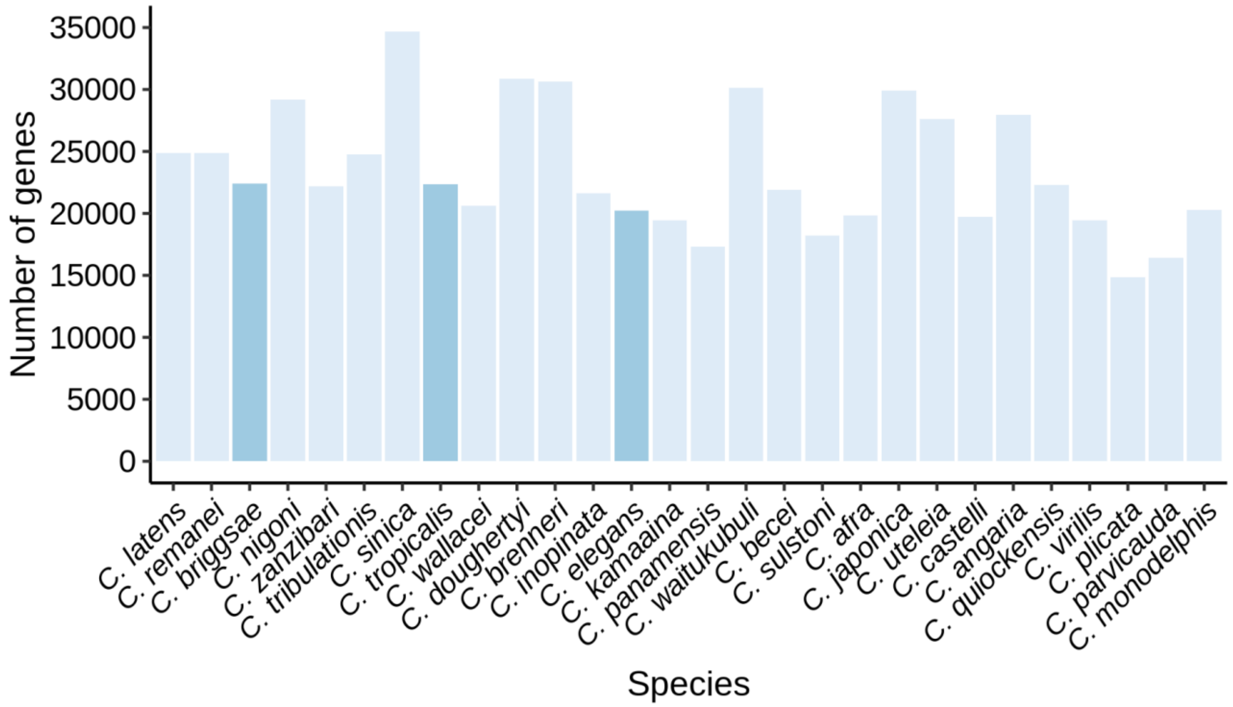


Supplemental Figure 2. Number of protein coding genes in the genomes used in this study. Species are highlighted by reproductive mode (light blue, female/male; dark blue, hermaphroditic).

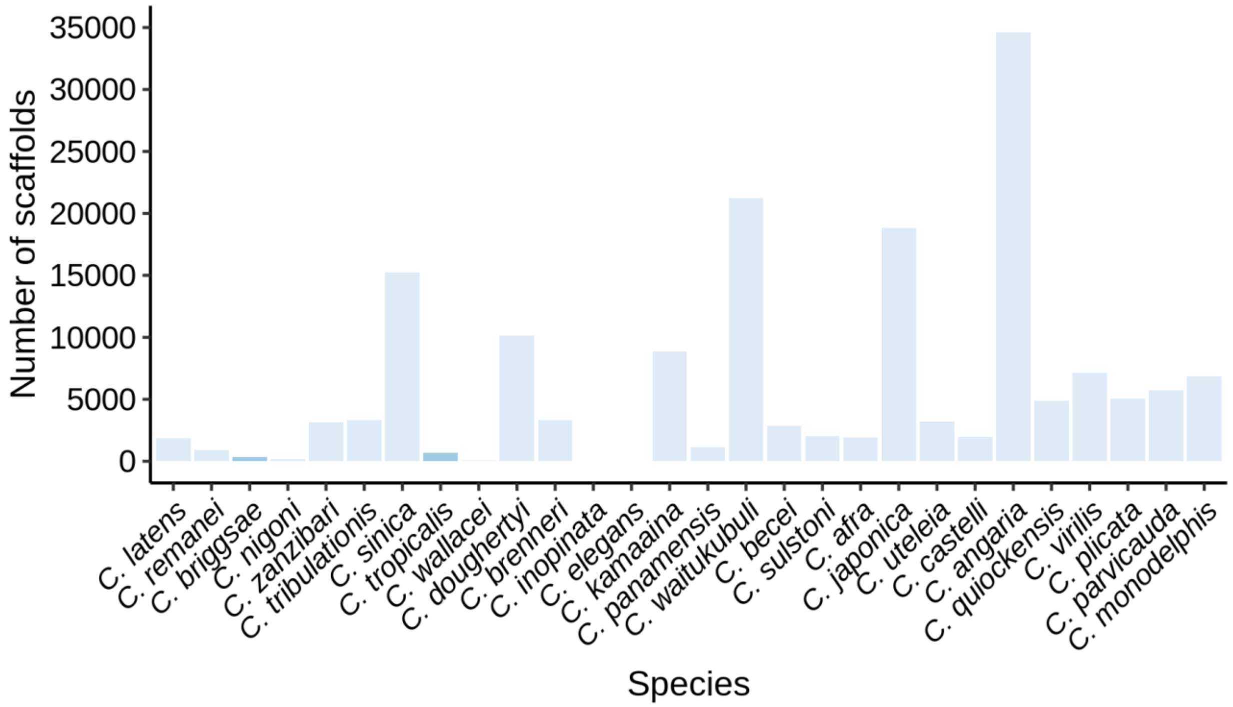


Supplemental Figure 3. Scaffold number among the genomes used in this study. Species are highlighted by reproductive mode (light blue, female/male; dark blue, hermaphroditic).

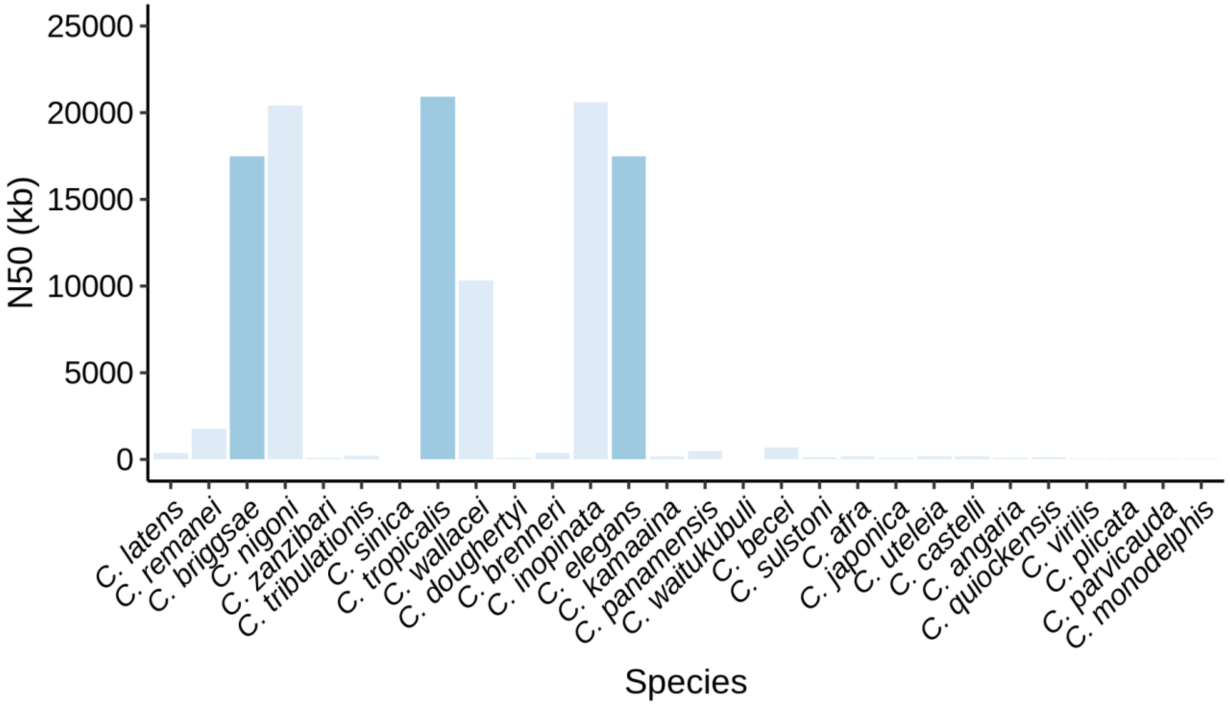


Supplemental Figure 4. N50 among the genomes used in this study. Species are highlighted by reproductive mode (light blue, female/male; dark blue, hermaphroditic).

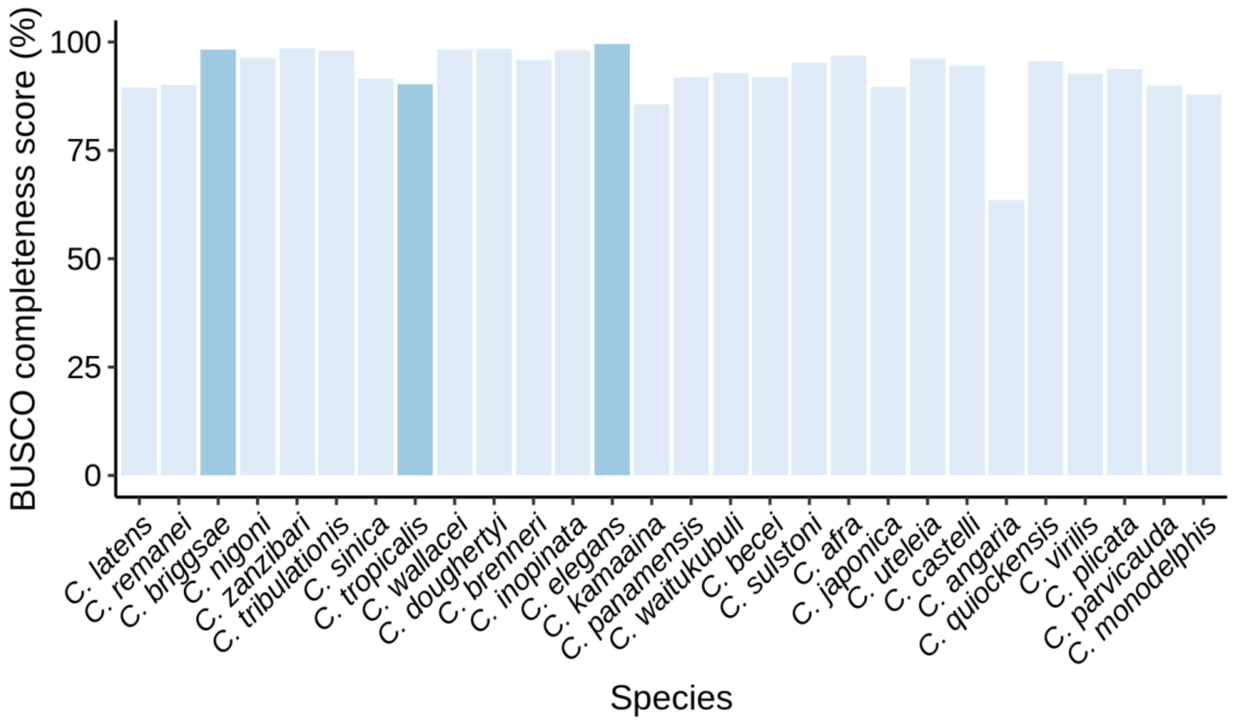


Supplemental Figure 5. BUSCO completeness score among the protein sets used in this study. Species are highlighted by reproductive mode (light blue, female/male; dark blue, hermaphroditic).

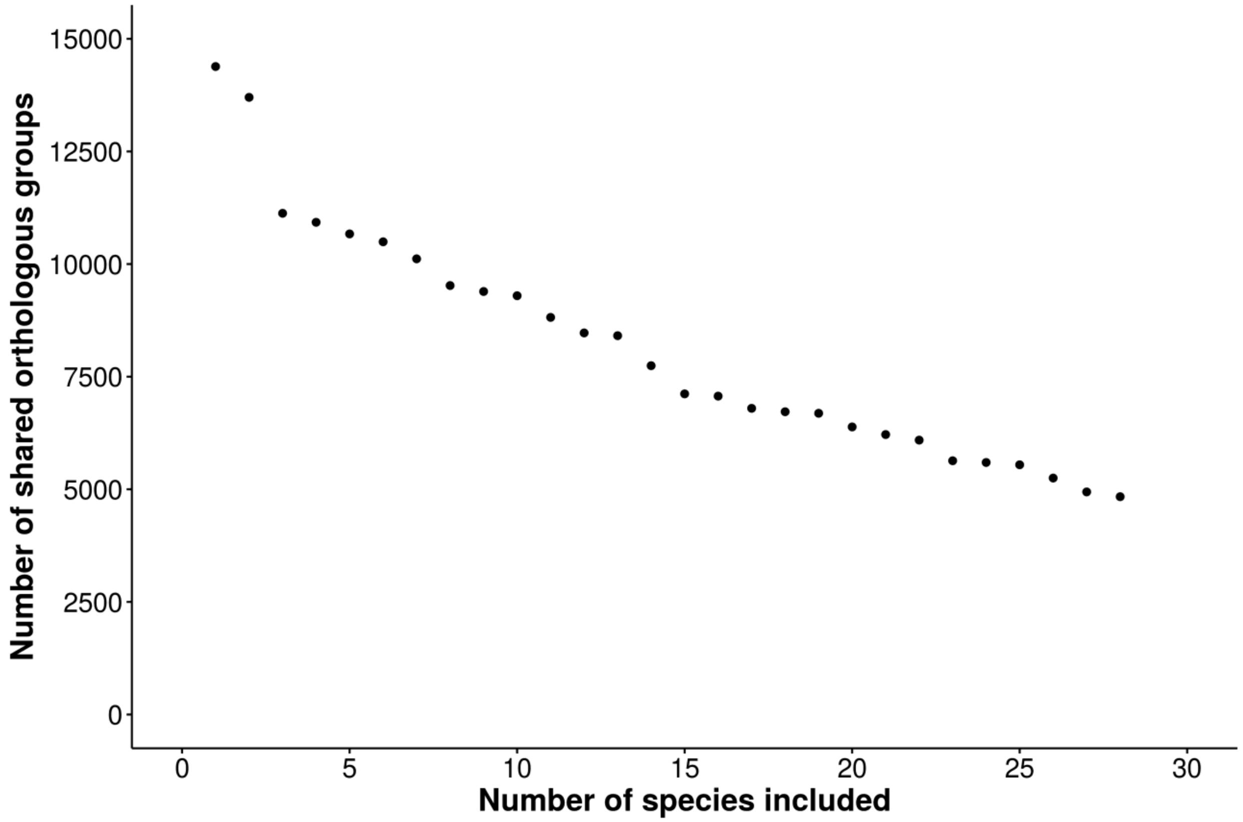


Supplemental Figure 6. The number of recovered shared orthologous groups decreases as more species are included. The first point is the number of orthologous groups recovered in *C. remanei*; the second point is the number of orthologous groups shared by *C. remanei* and *C. latens*. Subsequent points are the number of shared orthologous groups as more species are added in a sequence corresponding roughly to phylogenetic relatedness.

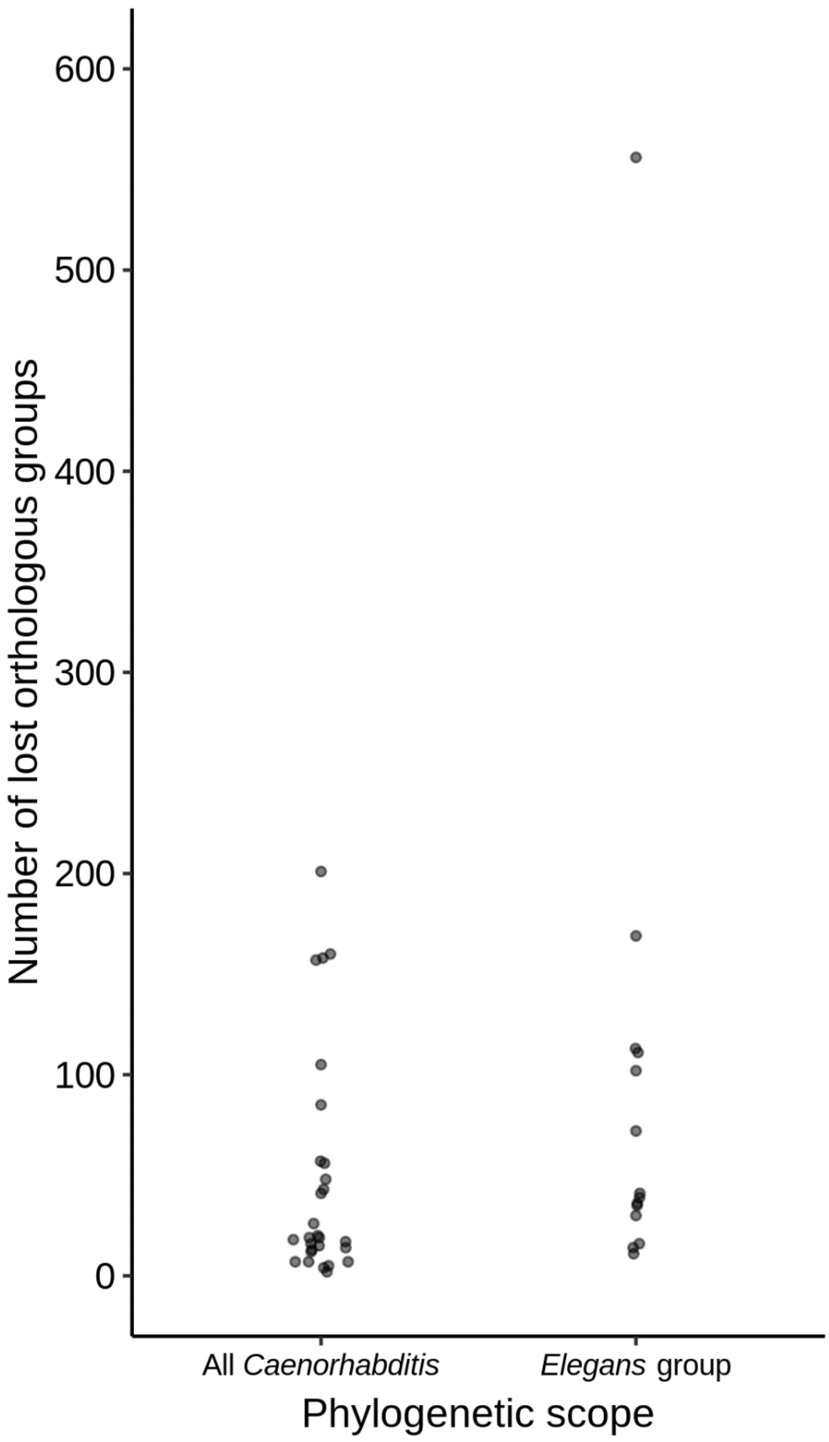


Supplemental Figure 7. The distribution of species-specific lost orthologous groups among all *Caenorhabditis* and only the *Elegans* group.

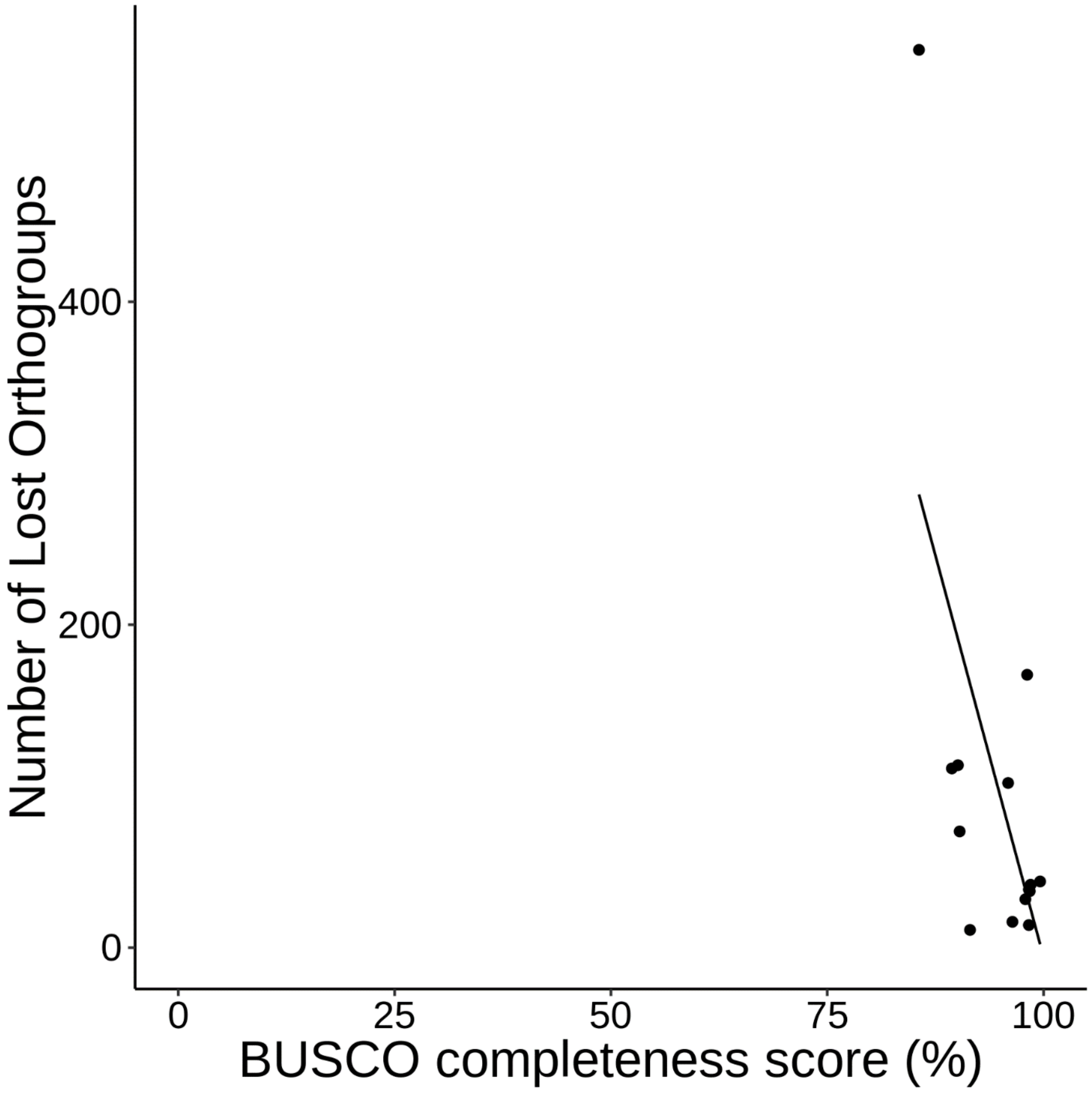


Supplemental Figure 8. The relationship between BUSCO completeness score and the number of lost orthologous groups when considering the *Elegans* group ($r^2$=0.36; *F*=8.3; β= -19.9; *p*=0.014).

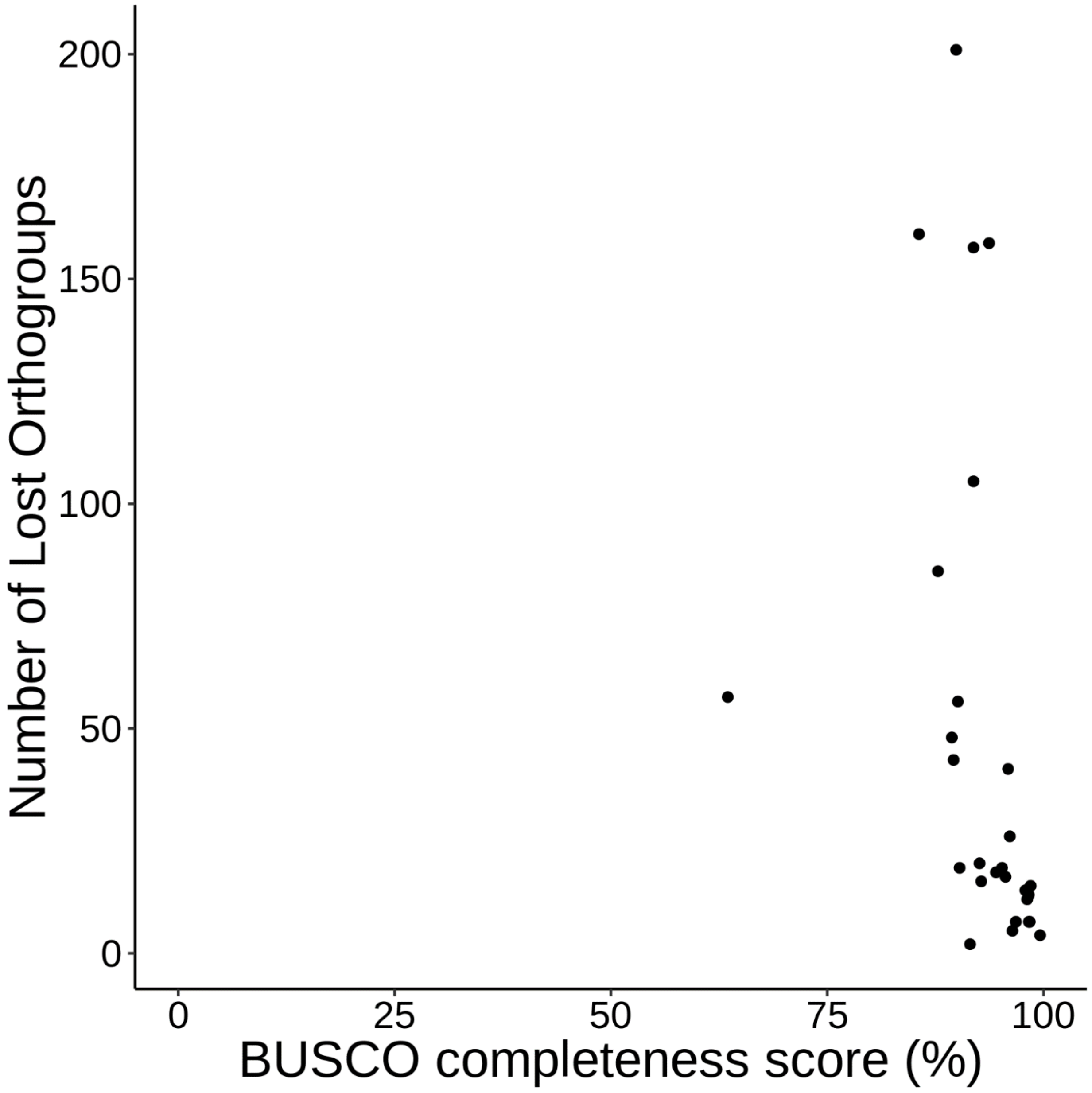


Supplemental Figure 9. The relationship between BUSCO completeness score and the number of lost orthologous groups when considering all *Caenorhabditis* ($r^2$=0.093; $F$=3.8; β= -2.9; $p$=0.063).

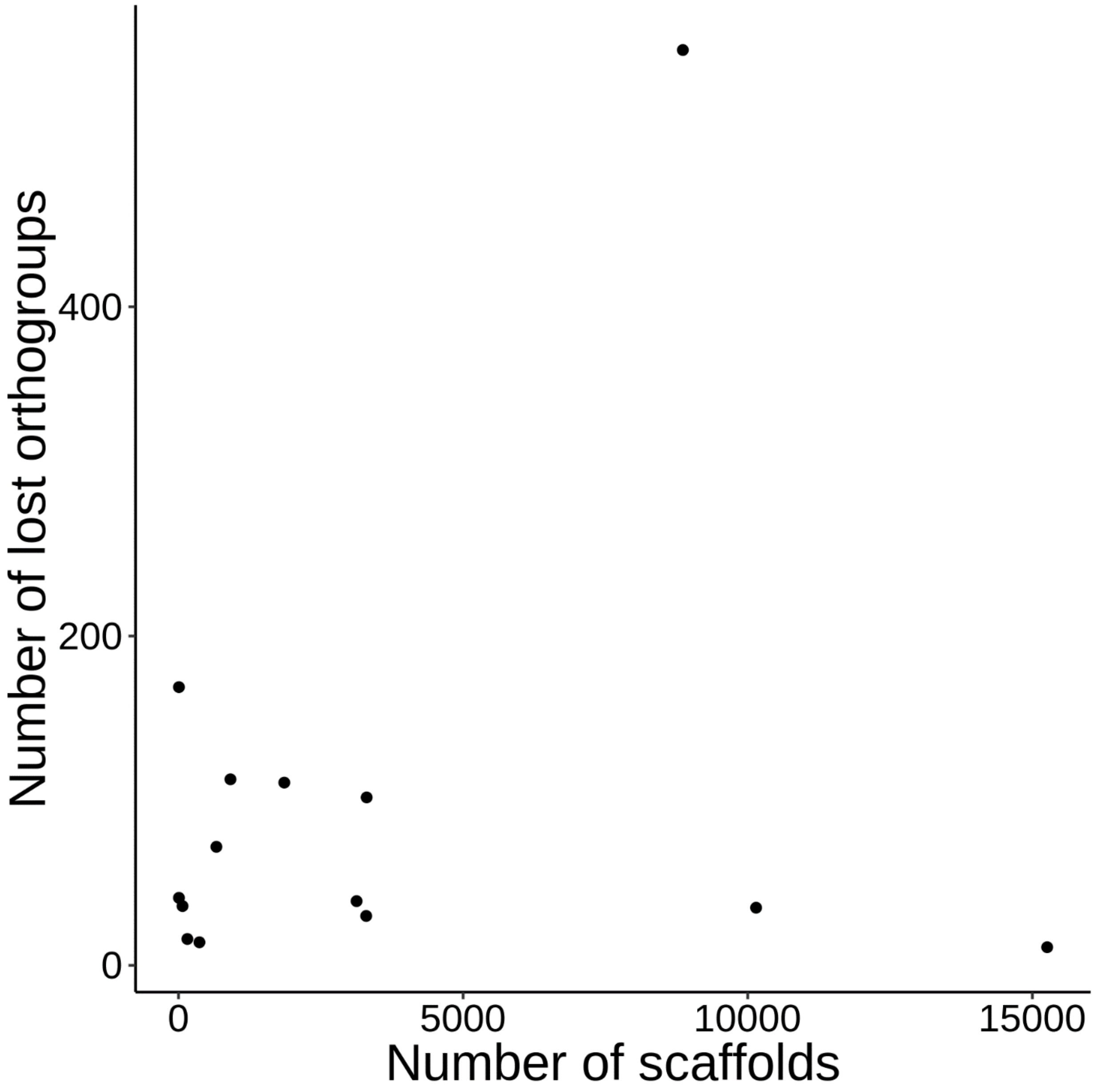


Supplemental Figure 10. The relationship between scaffold number and the number of lost orthologous groups when considering the *Elegans* group ($r^2$=0.038; $F$=0.53; β=0.0062; $p$=0.48).

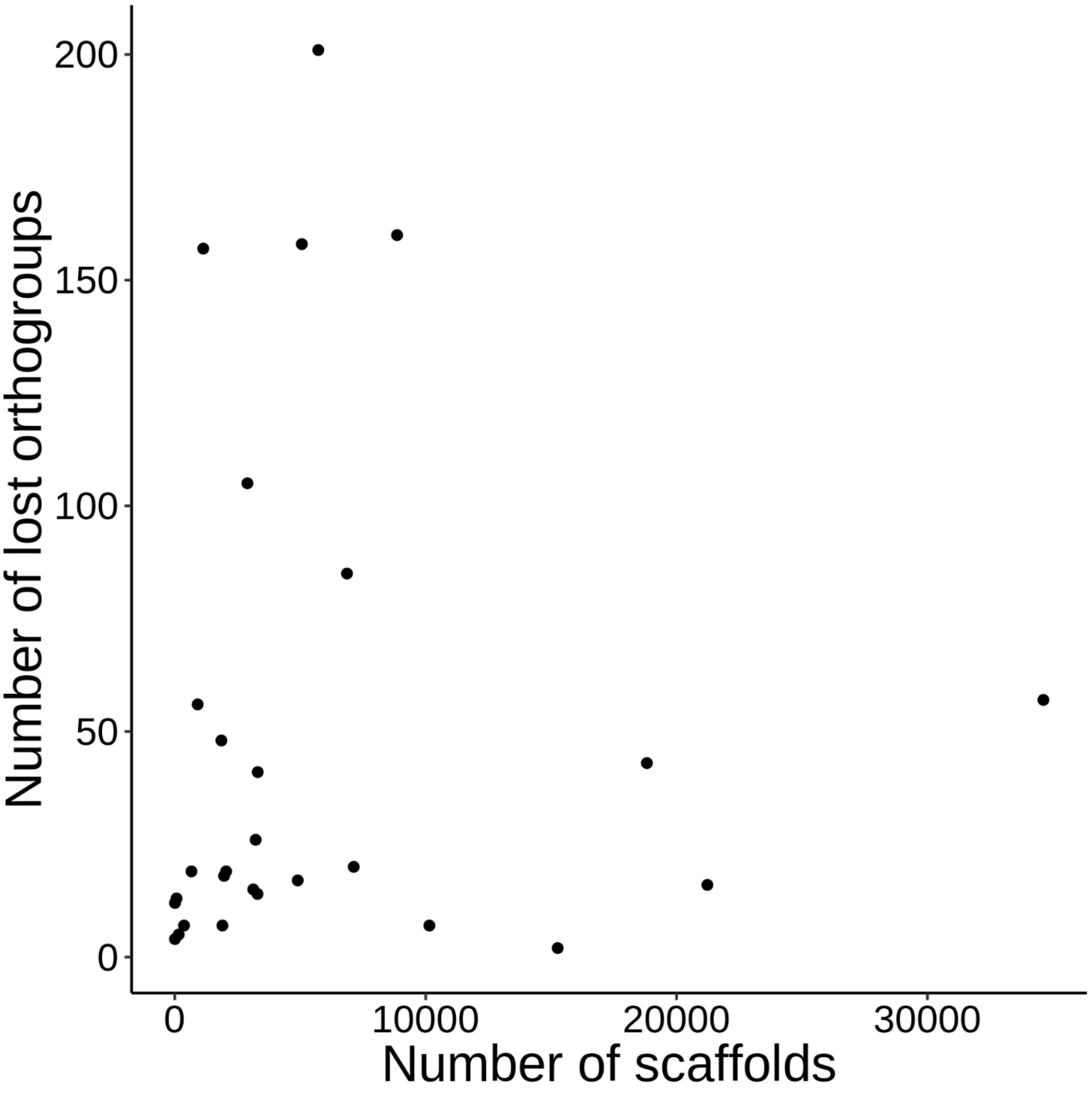


Supplemental Figure 11. The relationship between scaffold number and the number of lost orthologous groups when considering all *Caenorhabditis* ($r^2$=0.036; $F$=0.057; β=0.00033; $p$=0.81).

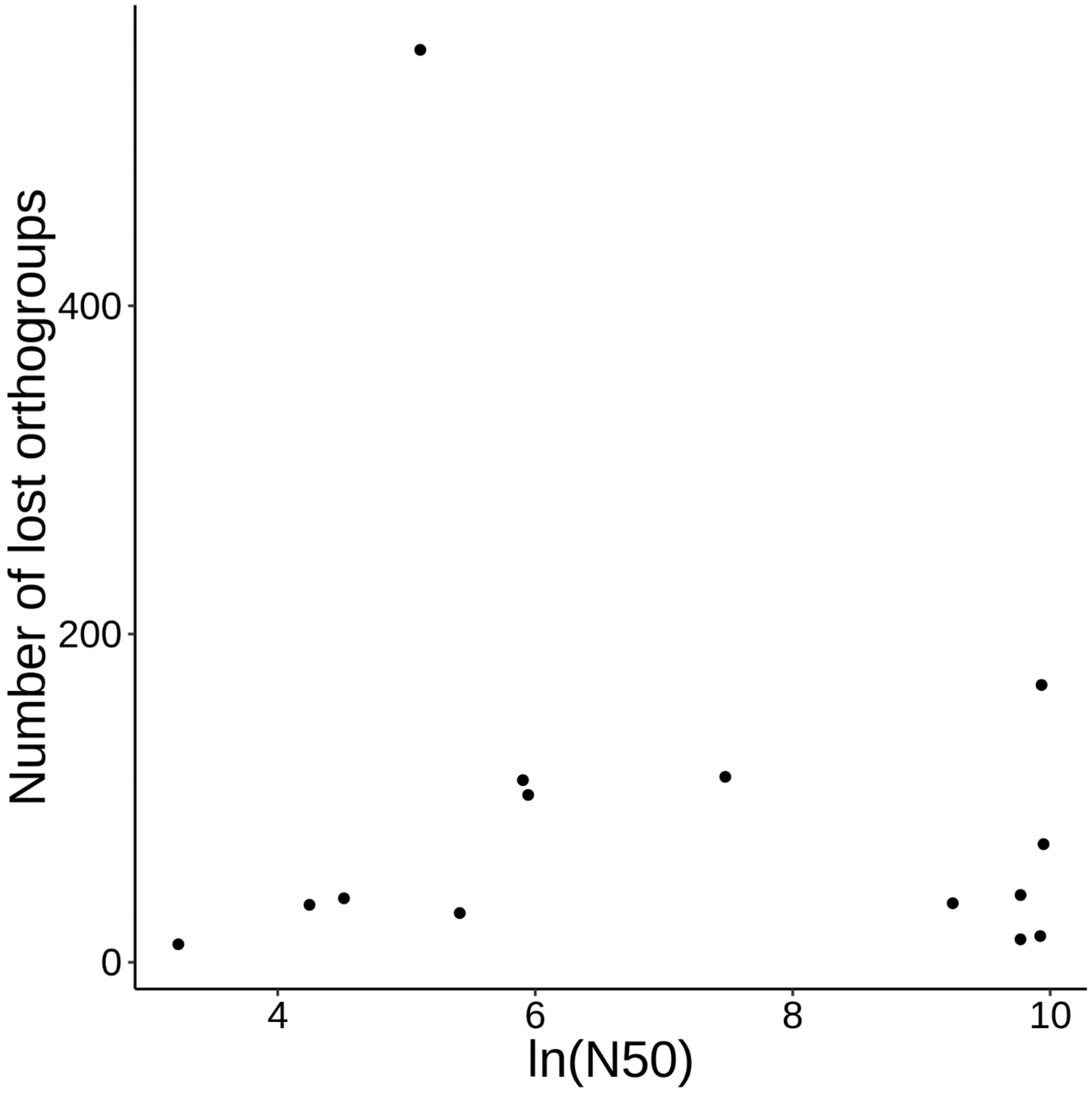


Supplemental Figure 12. The relationship between N50 (here, ln(N50)) and the number of lost orthologous groups when considering the *Elegans* group ($r^2$=0.054; $F$=0.33; β= -9.2; $p$=0.58).

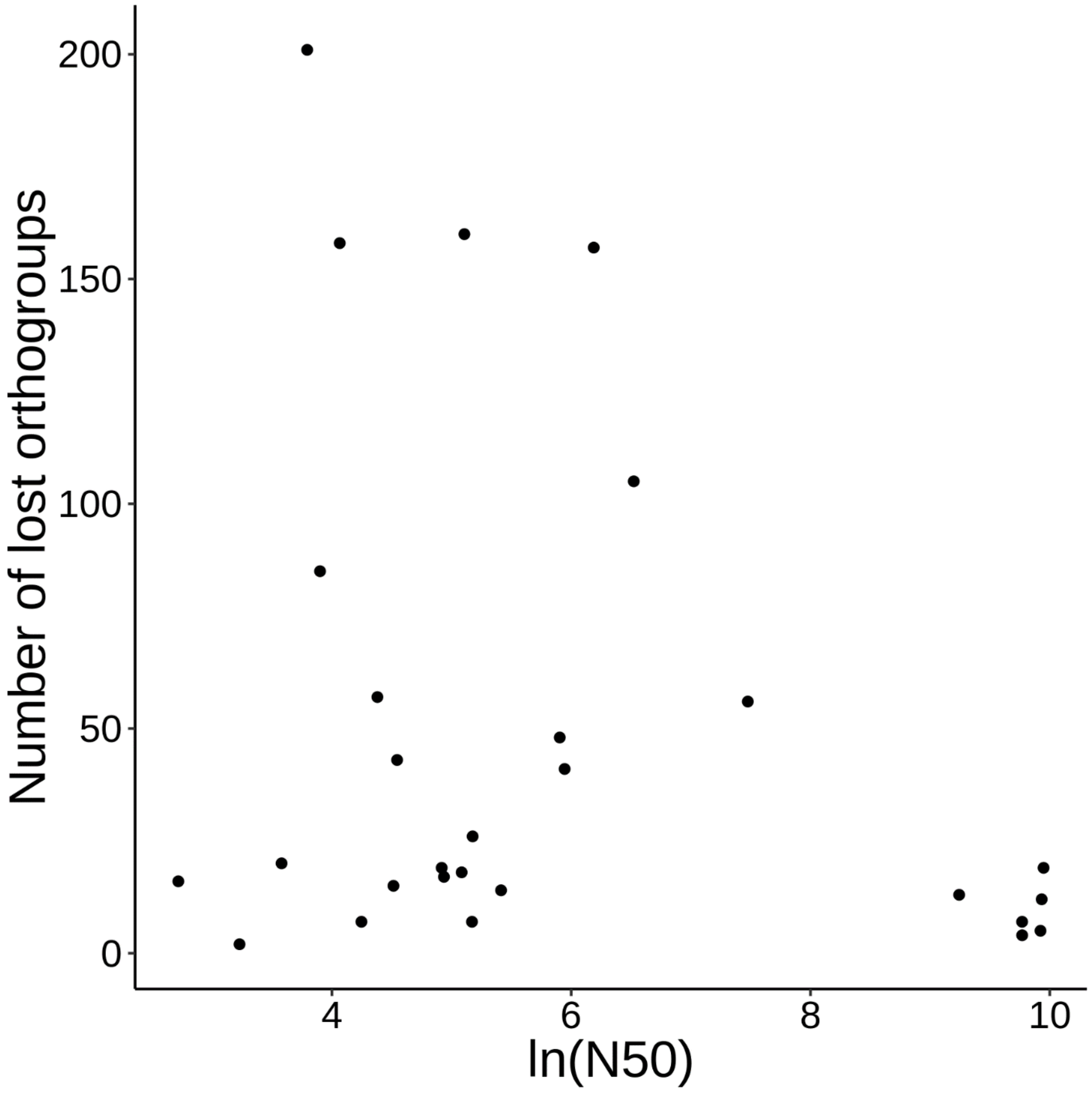


Supplemental Figure 13. The relationship between N50 (here, ln(N50)) and the number of lost orthologous groups when considering all *Caenorhabditis* ($r^2$=0.038; $F$=2.1; β= -6.7; $p$=0.16).

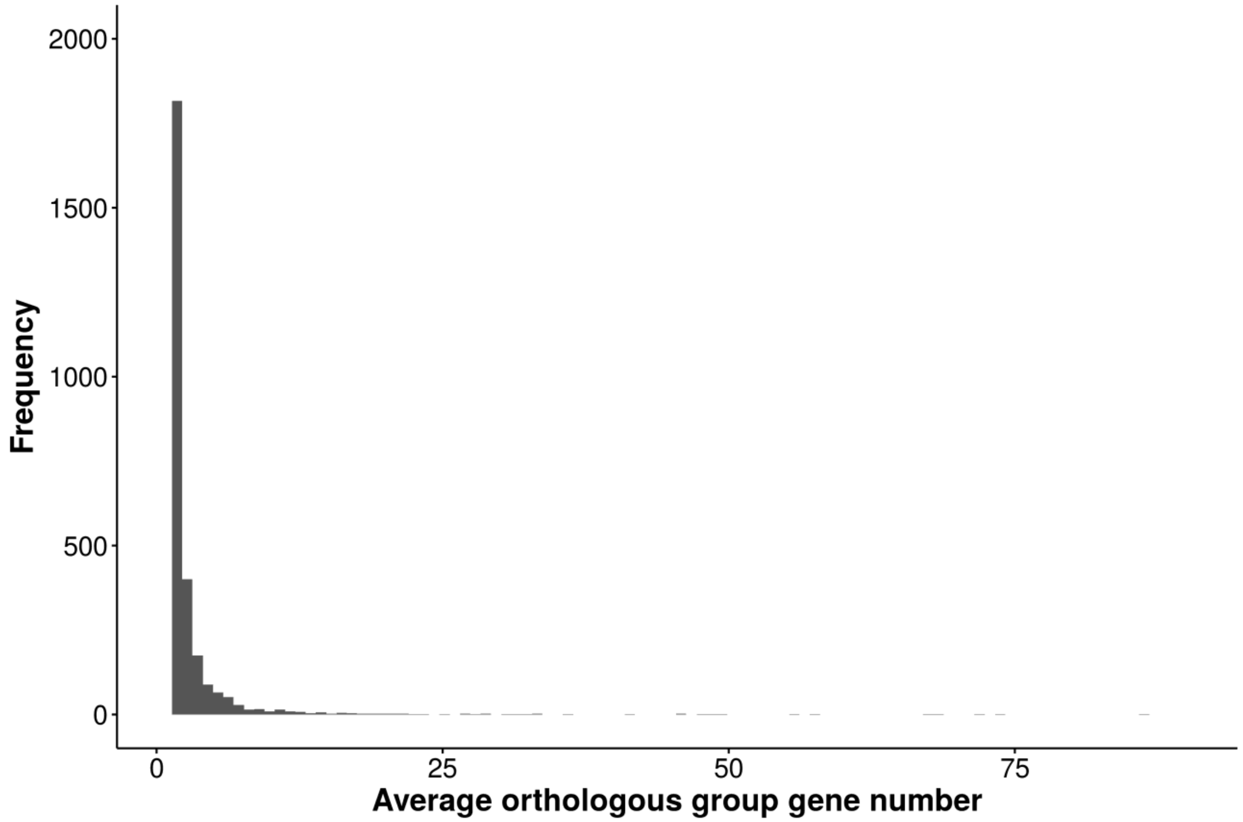


Supplemental Figure 14. The average gene copy number per orthologous group is low (mean=0.81; range 0.07-86.0). Here, the y-axis has been arbitrarily cut off at 2000 for readability. Species-specific orthologous groups are not included here.

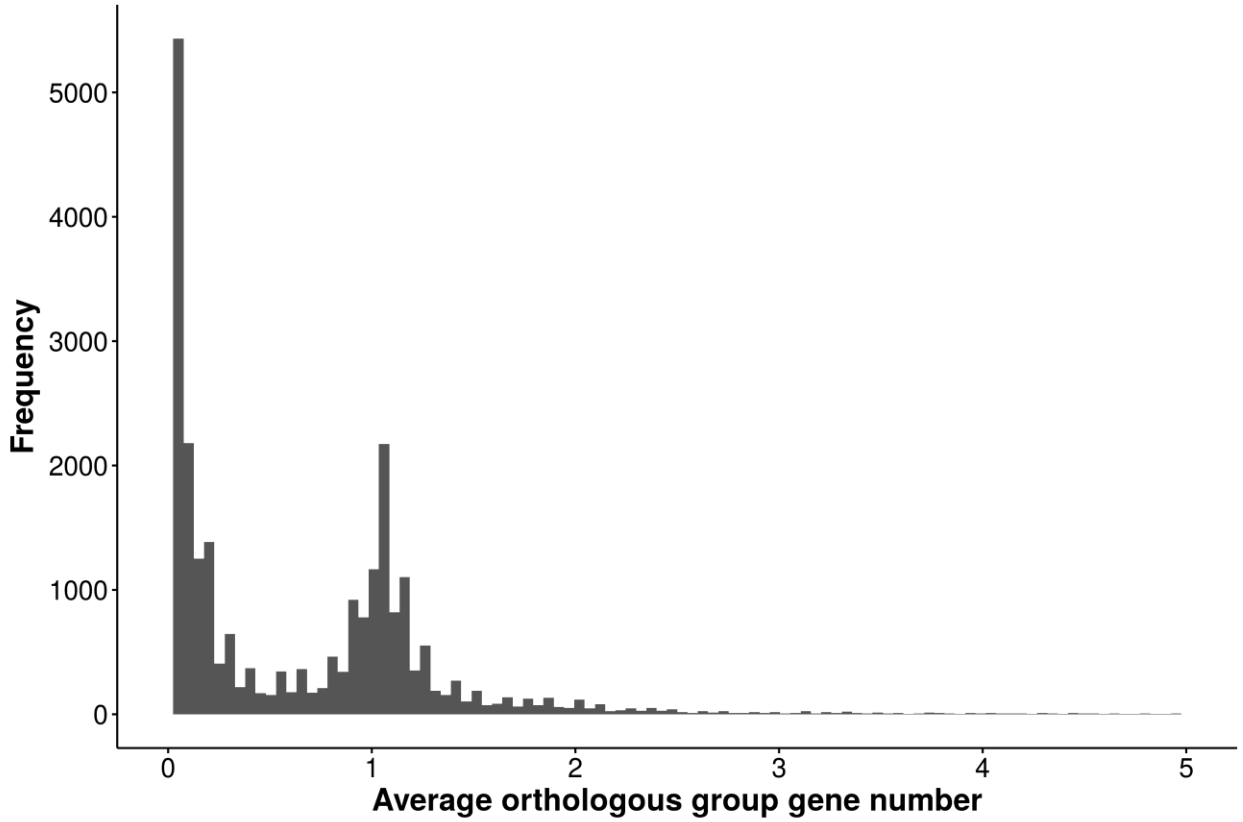


Supplemental Figure 15. The average gene copy number per orthologous group is low. Here, only orthologous groups with average gene copy number less than five are plotted. Most defined orthologous groups are present in only a few species, and a second peak appears at orthologous groups that are nearly single-copy across all species.

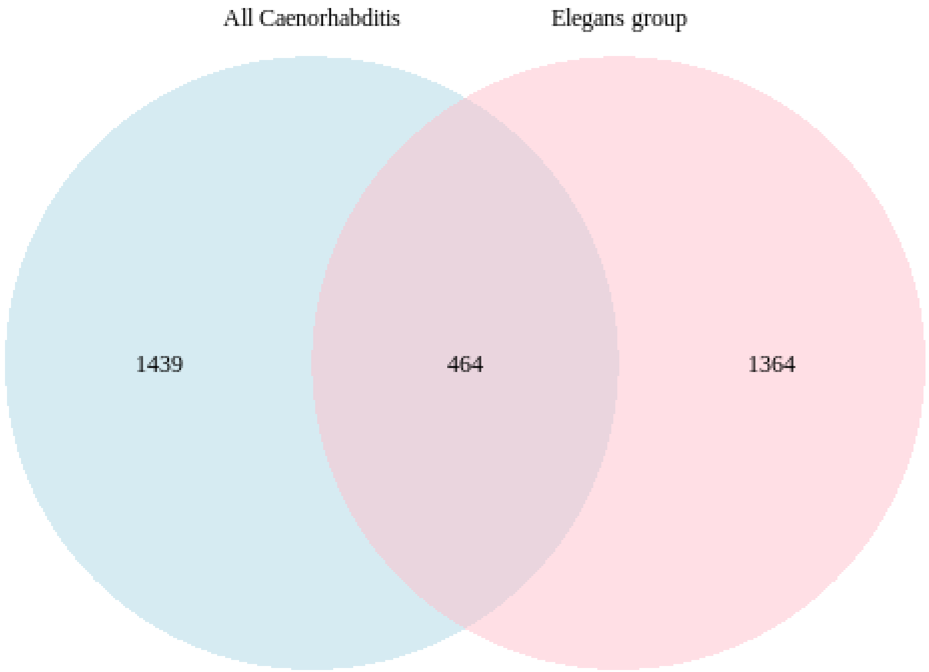


Supplemental Figure 16. The *C. elegans* gene constituents of species-specific lost orthologous groups do not largely overlap among different levels of phylogenetic consideration (all *Caenorhabditis* or the *Elegans* group).

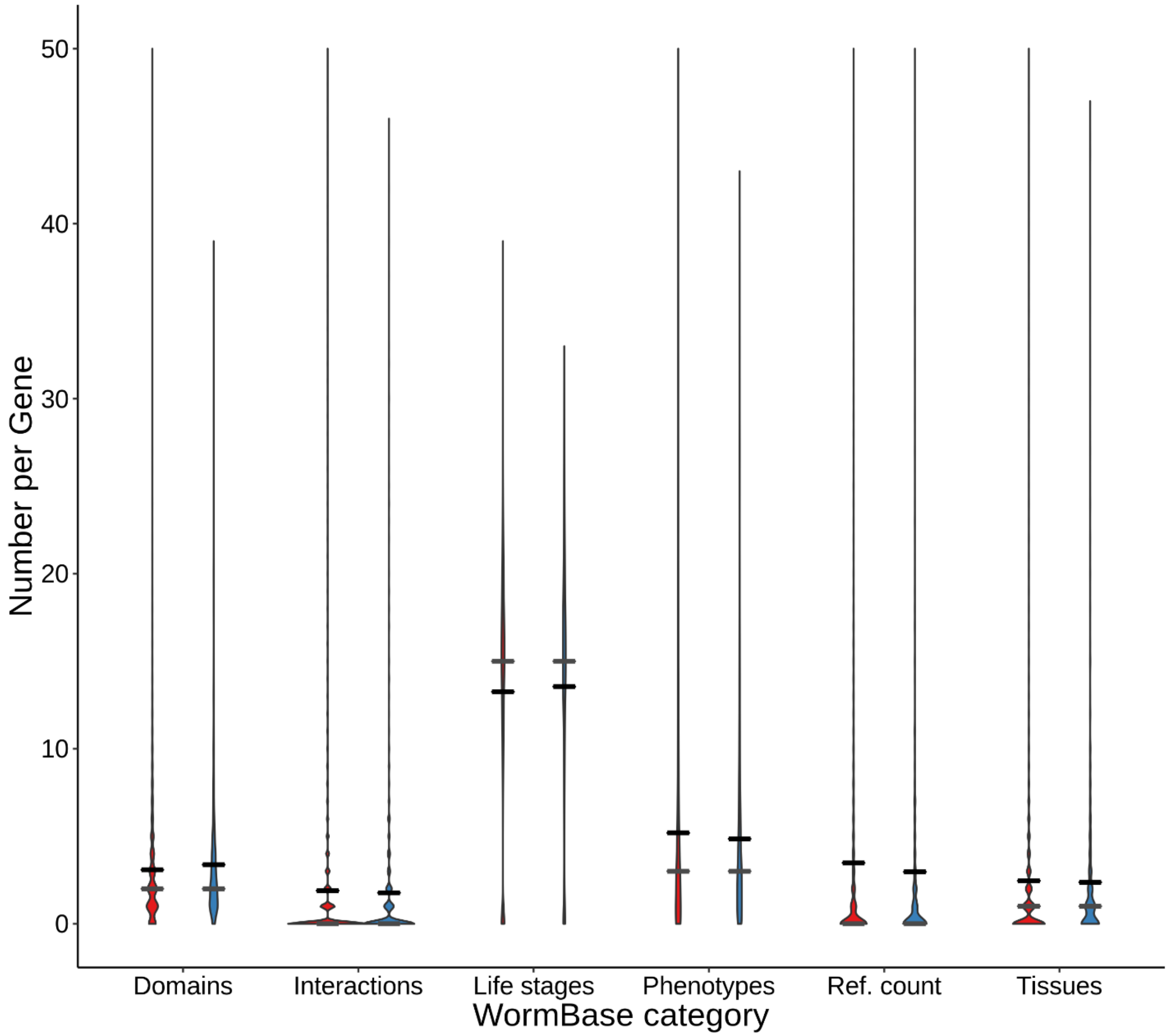


Supplemental Figure 17. The distribution of number of unique WormBase terms or Pfam domains among retained (red) and lost (blue) genes when considering all *Caenorhabditis*. Black bars, means. Gray bars, medians. Here, the y-axis has been arbitrarily limited to 50 to make the figure readable. This excludes 585 data points (out of 118296) from the violin plots, but all data is used for plotting means and medians.

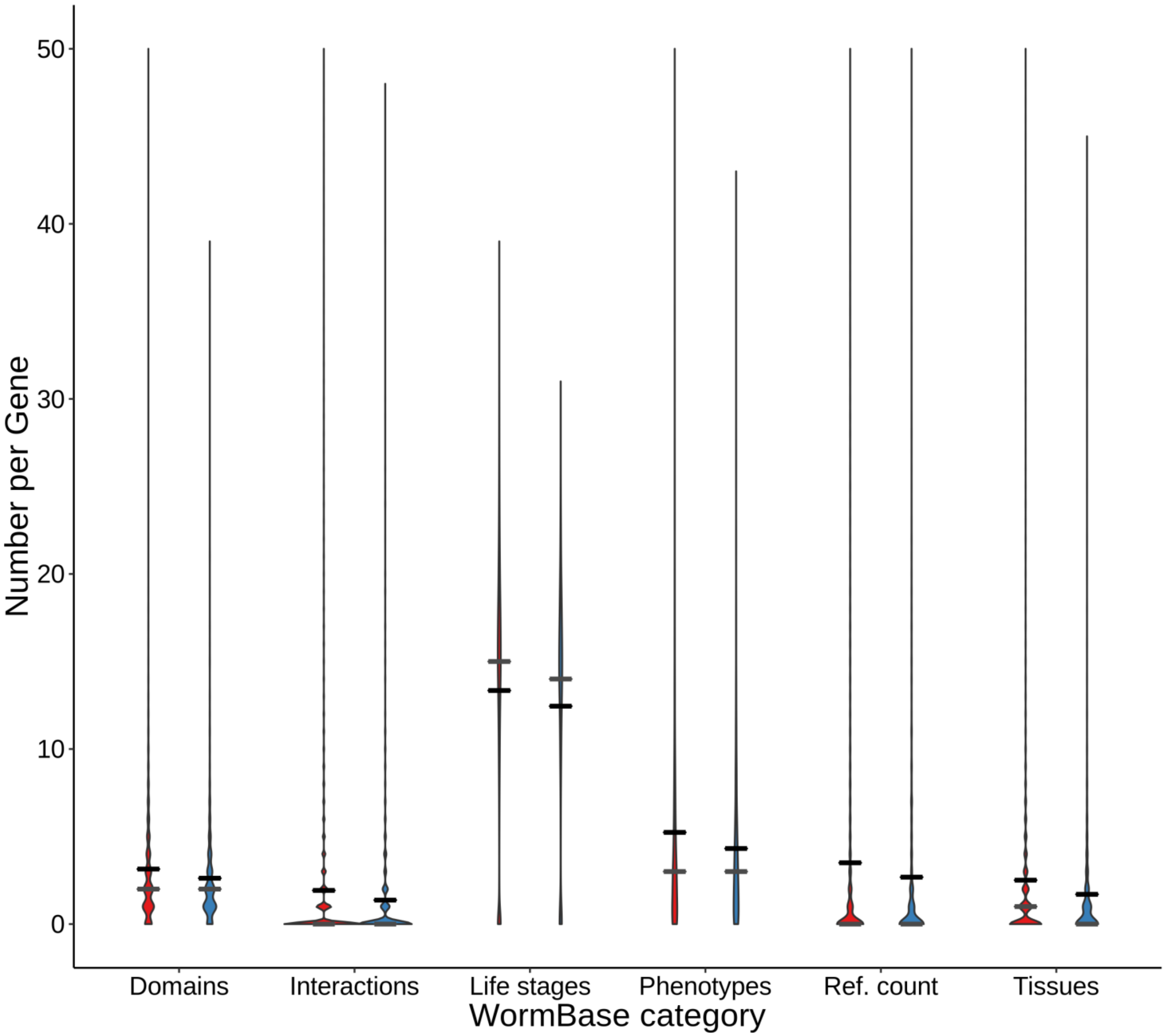


Supplemental Figure 18. The distribution of number of unique WormBase terms or Pfam domains among retained (red) and lost (blue) genes when considering the *Elegans* group. Black bars, means. Gray bars, medians. Here, the y-axis has been arbitrarily limited to 50 to make the figure readable. This excludes 585 data points (out of 118296) from the violin plots, but all data is used for plotting means and medians.

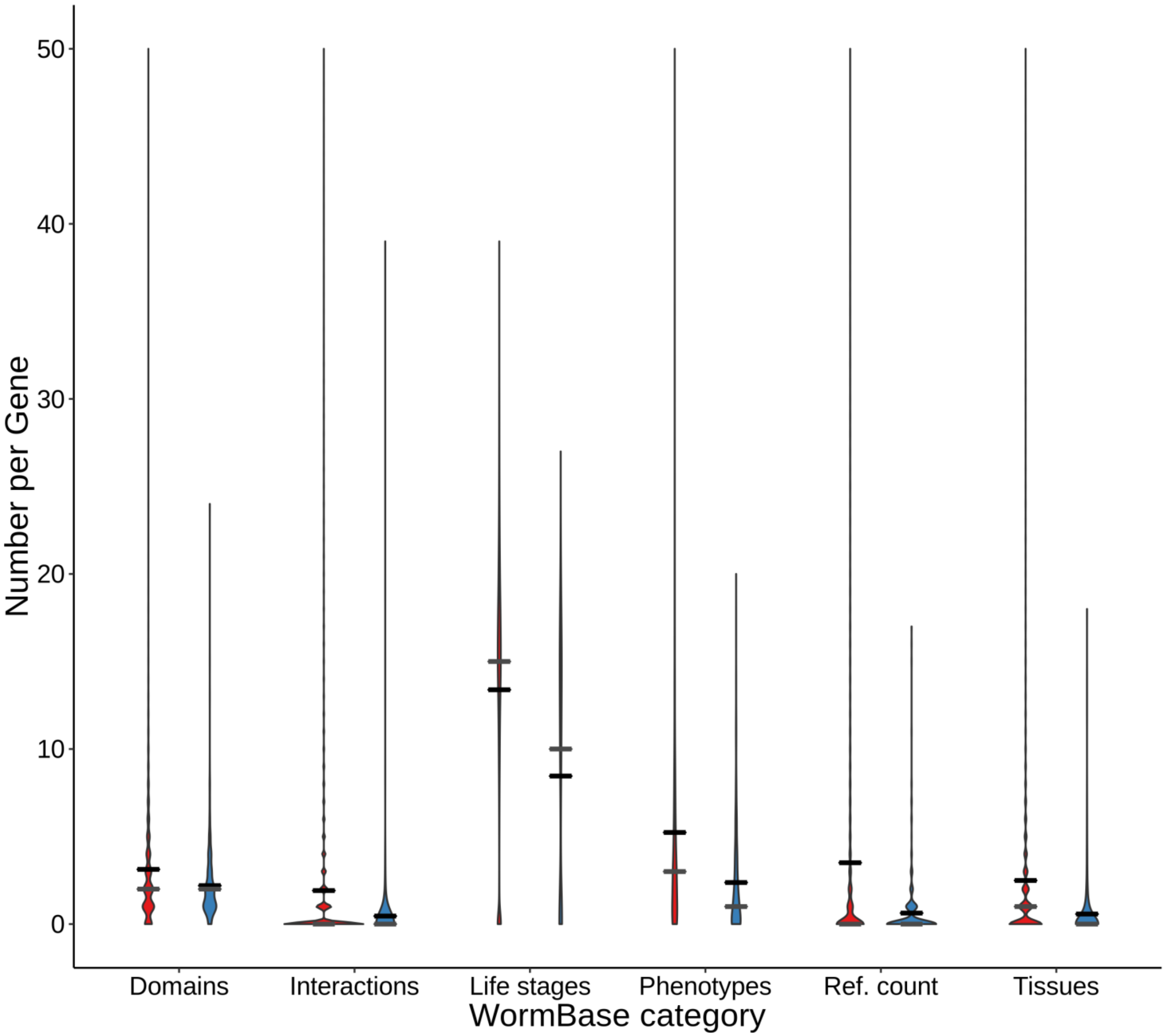


Supplemental Figure 19. The distribution of number of unique WormBase terms or Pfam domains among retained (red) and lost (blue) genes when considering only *C. inopinata*. Black bars, means. Gray bars, medians. Here, the y-axis has been arbitrarily limited to 50 to make the figure readable. This excludes 585 data points (out of 118296) from the violin plots, but all data is used for plotting means and medians

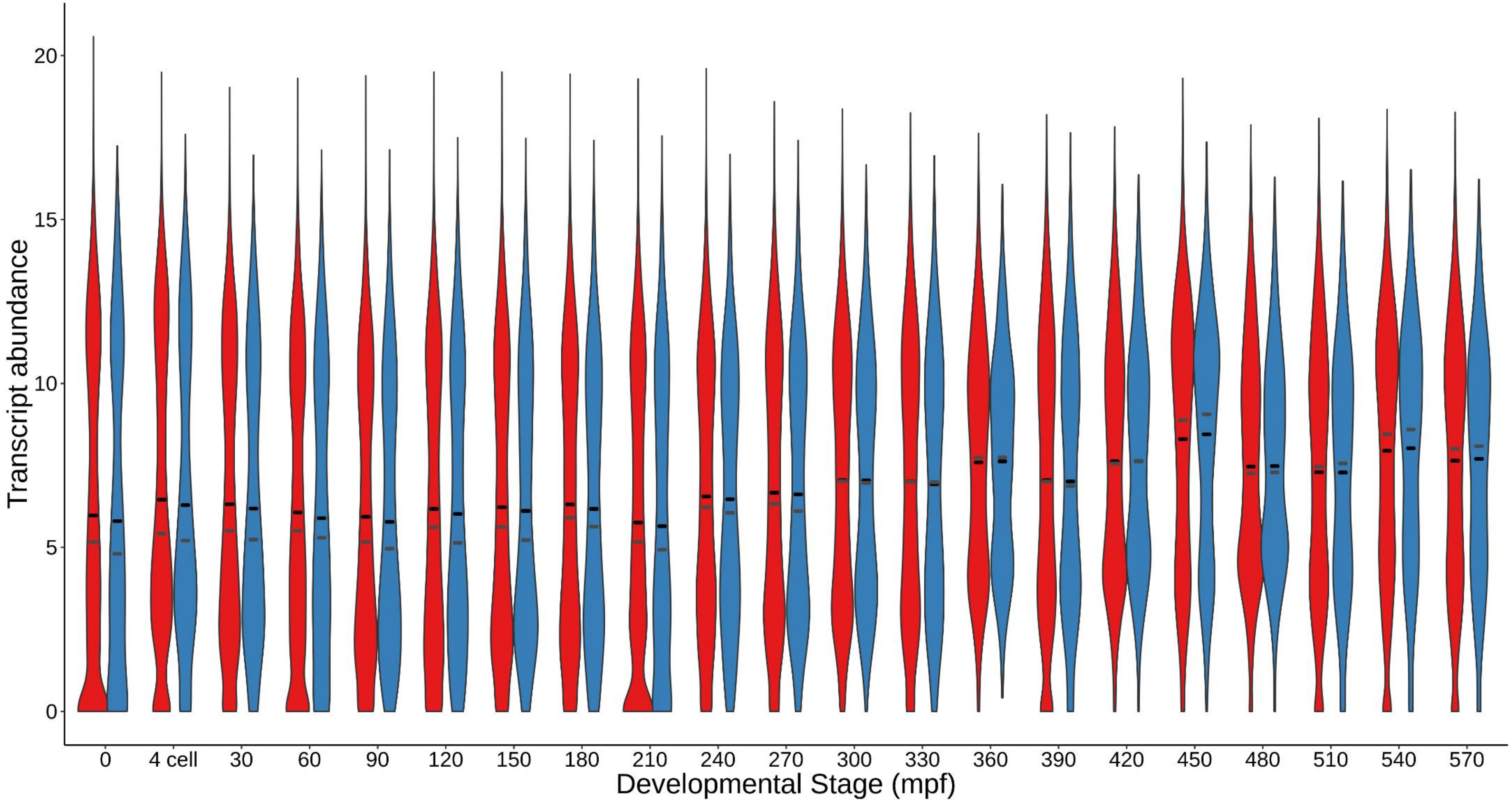


Supplemental Figure 20. The distribution of transcriptional abundance by embryonic stage among retained (red) and lost (blue) genes when considering all *Caenorhabditis*. Black bars, means. Gray bars, medians. Here, the transcriptional activity is measured in units of $log_2$(1+dpcm) on the y-axis. Data were originally reported in Boeck et al. 2016.

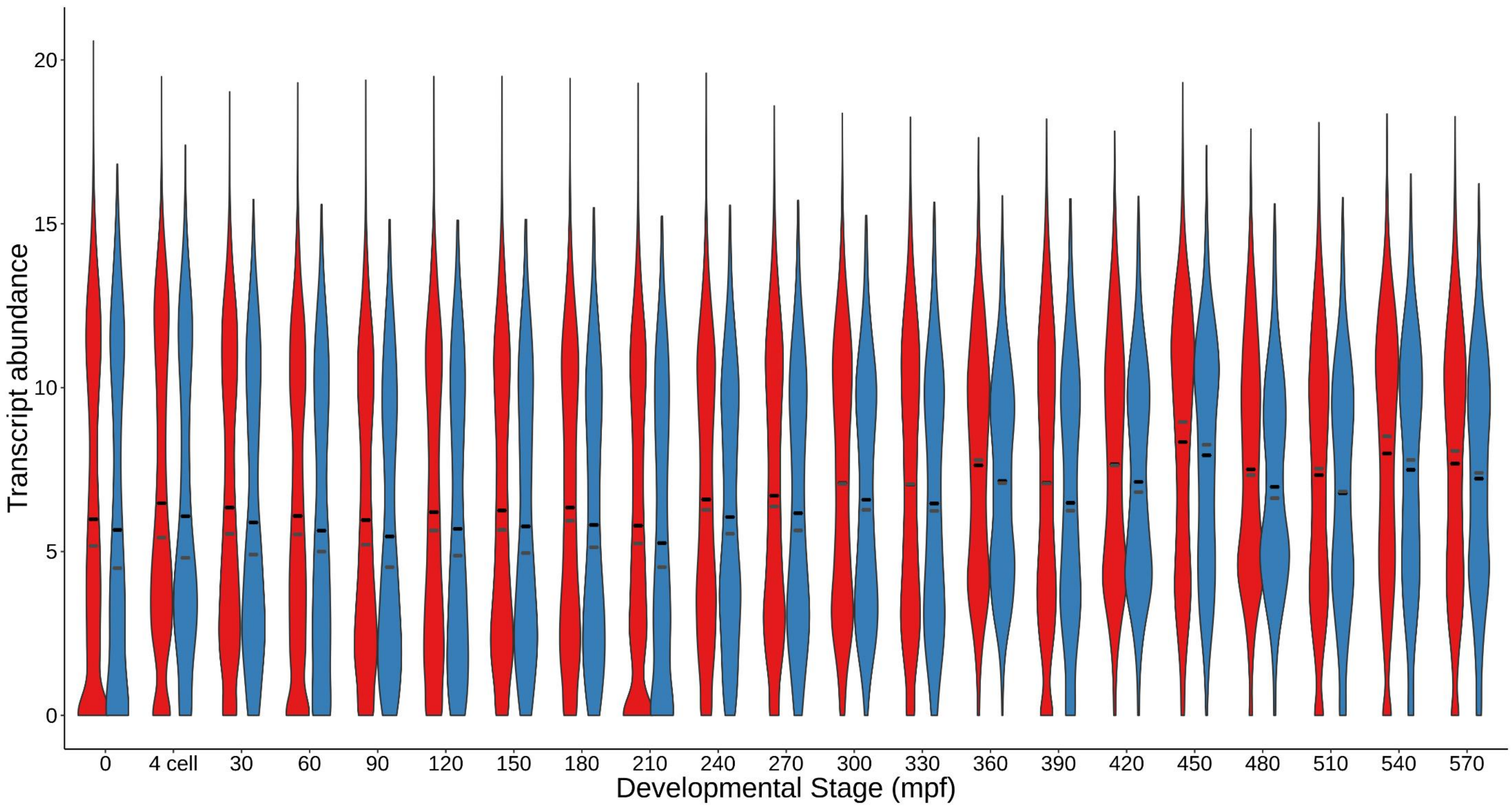


Supplemental Figure 21. The distribution of transcriptional abundance by embryonic stage among retained (red) and lost (blue) genes when considering the *Elegans* group. Black bars, means. Gray bars, medians. Here, the transcriptional activity is measured in units of $\log_2(1+\text{dpcm})$ on the y-axis. Data were originally reported in Boeck et al. 2016.

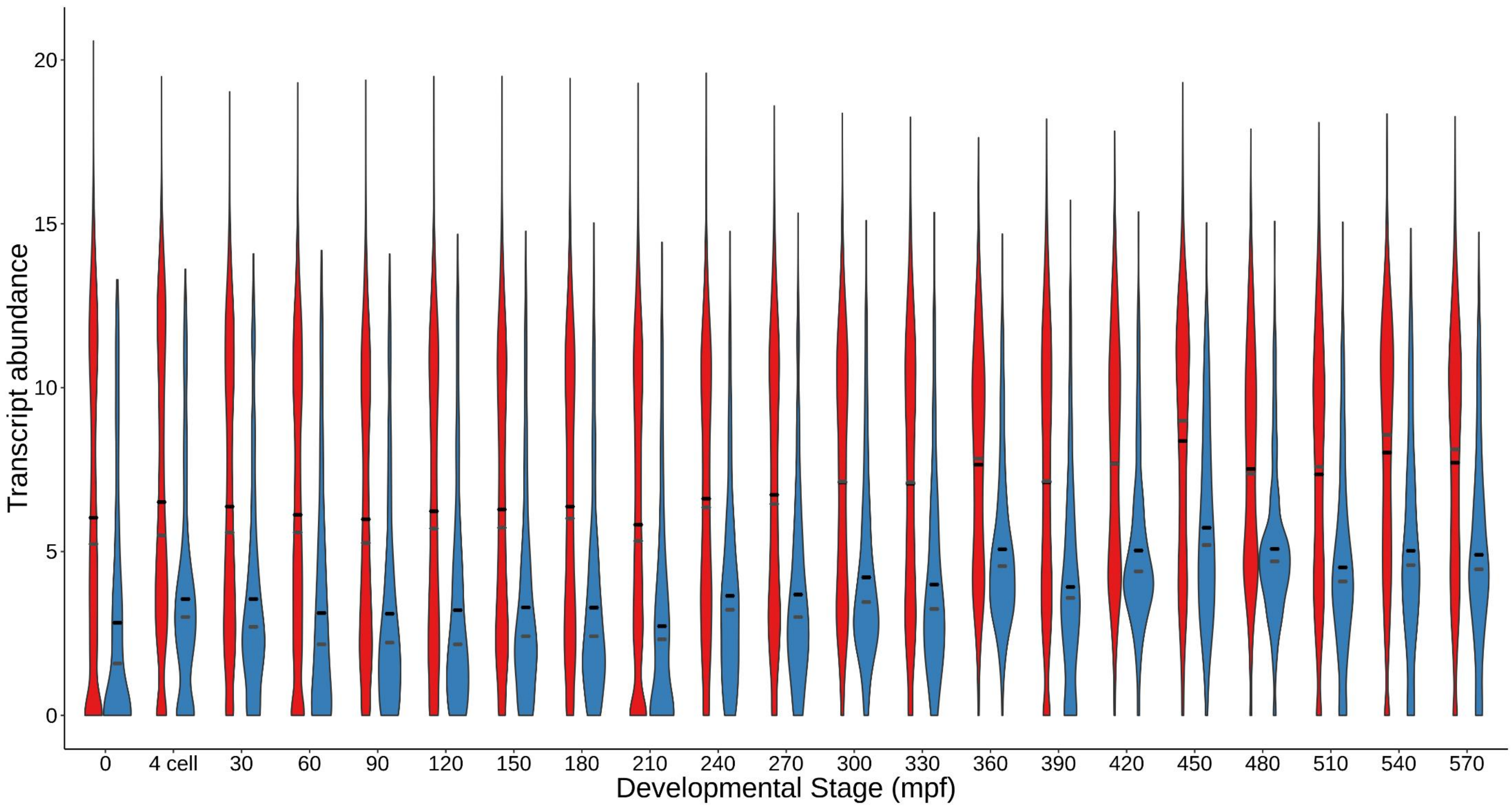


Supplemental Figure 22. The distribution of transcriptional abundance by embryonic stage among retained (red) and lost (blue) genes when considering only *C. inopinata*. Black bars, means. Gray bars, medians. Here, the transcriptional activity is measured in units of $\log_2(1+\text{dpcm})$ on the y-axis. Data were originally reported in Boeck et al. 2016.

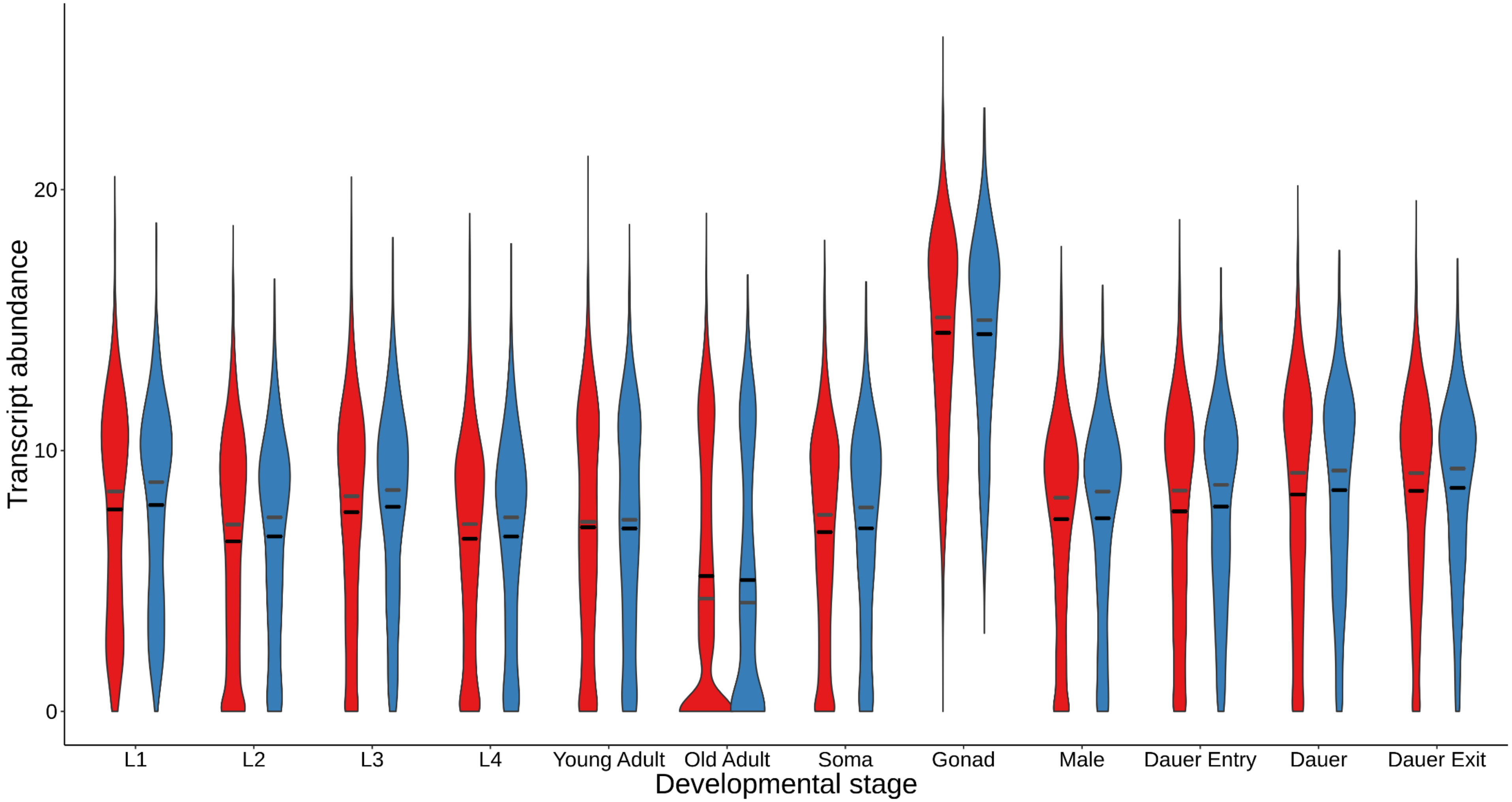


Supplemental Figure 23. The distribution of transcriptional abundance by postembryonic stage among retained (red) and lost (blue) genes when considering all *Caenorhabditis*. Black bars, means. Gray bars, medians. Here, the transcriptional activity is measured in units of $\log_2$(1+dpcm) on the y-axis. Data were originally reported in Boeck et al. 2016.

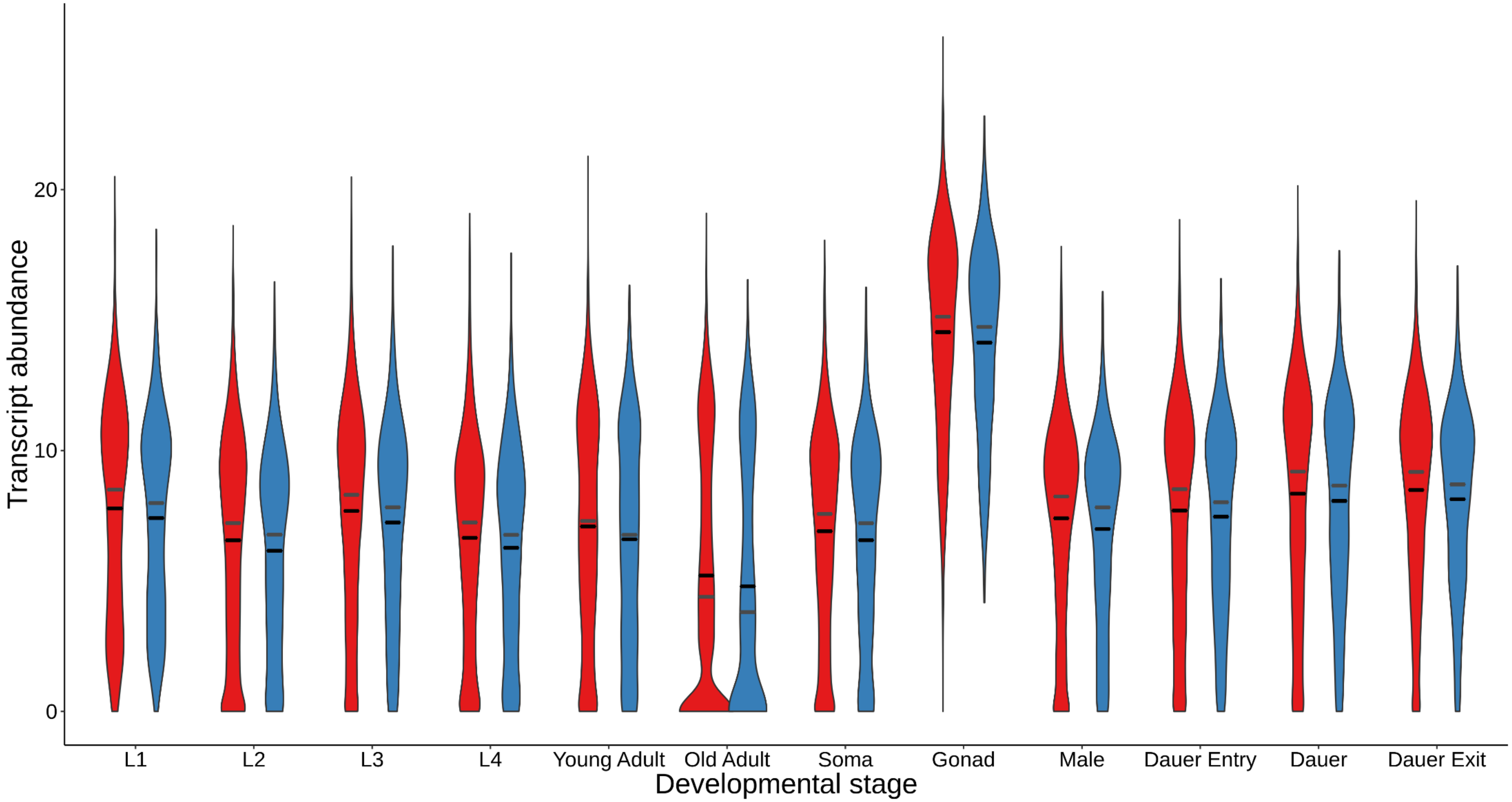


Supplemental Figure 24. The distribution of transcriptional abundance by postembryonic stage among retained (red) and lost (blue) genes when considering the *Elegans* group. Black bars, means. Gray bars, medians. Here, the transcriptional activity is measured in units of $\log_2(1+\text{dpcm})$ on the y-axis. Data were originally reported in Boeck et al. 2016.

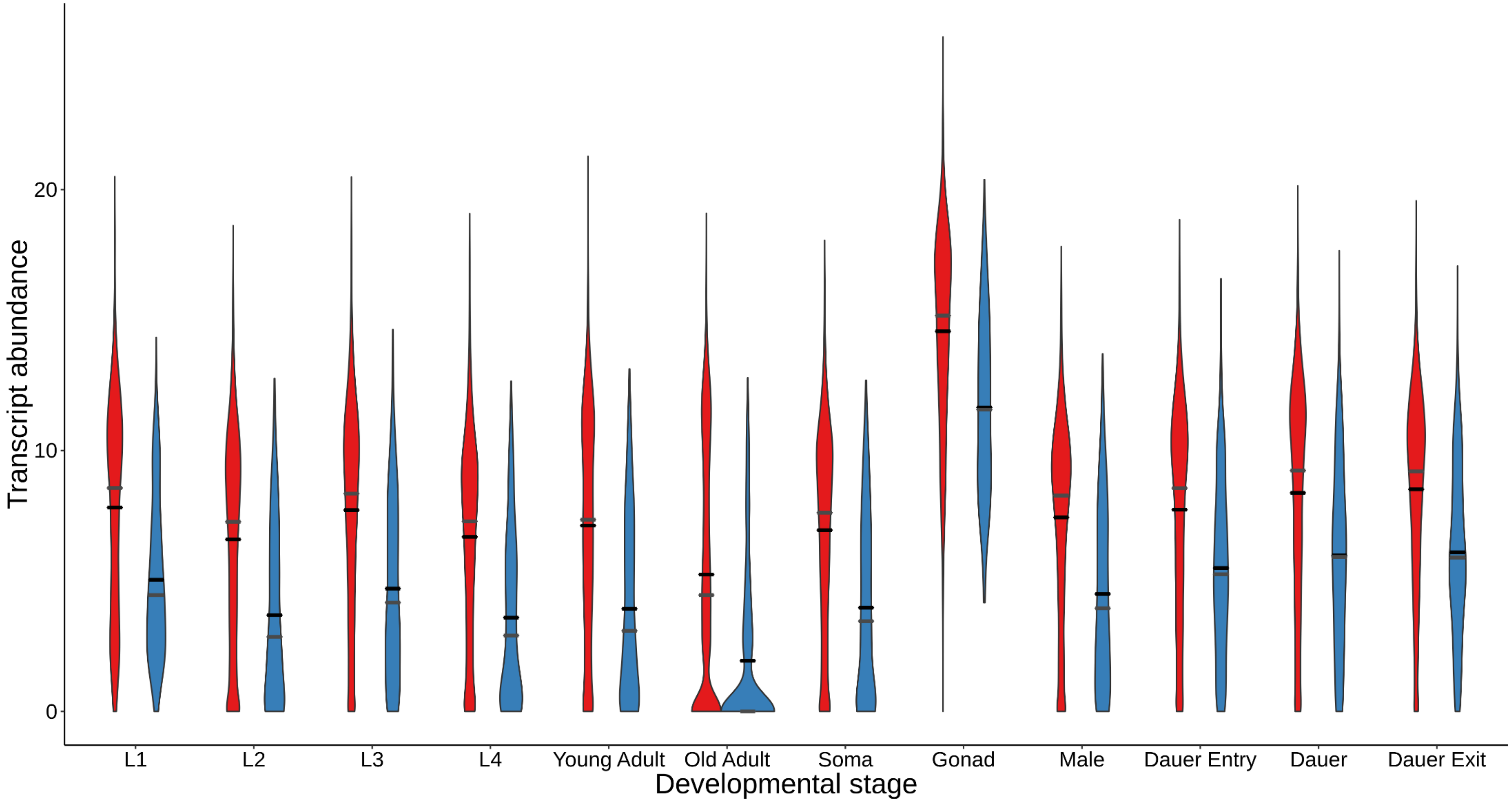


Supplemental Figure 25. The distribution of transcriptional abundance by postembryonic stage among retained (red) and lost (blue) genes when considering only *C. inopinata*. Black bars, means. Gray bars, medians. Here, the transcriptional activity is measured in units of $\log_2$(1+dpcm) on the y-axis. Data were originally reported in Boeck et al. 2016.

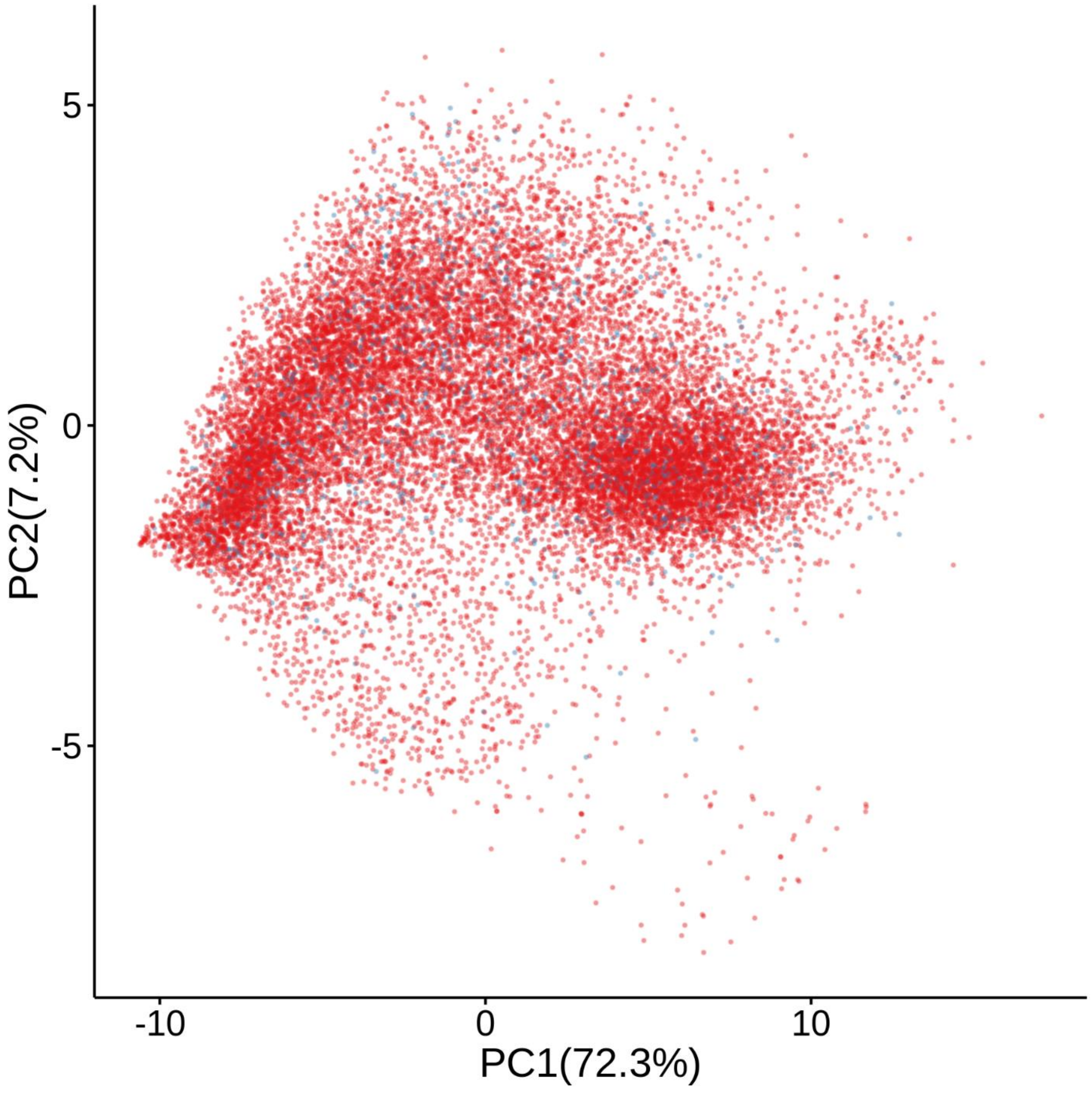


Supplemental Figure 26. Principal components analysis of transcriptomic and WormBase data. Points colored by loss when considering all *Caenorhabditis*. Lost genes do not appear to cluster in multidimensional space. Blue, lost genes. Red, retained genes.

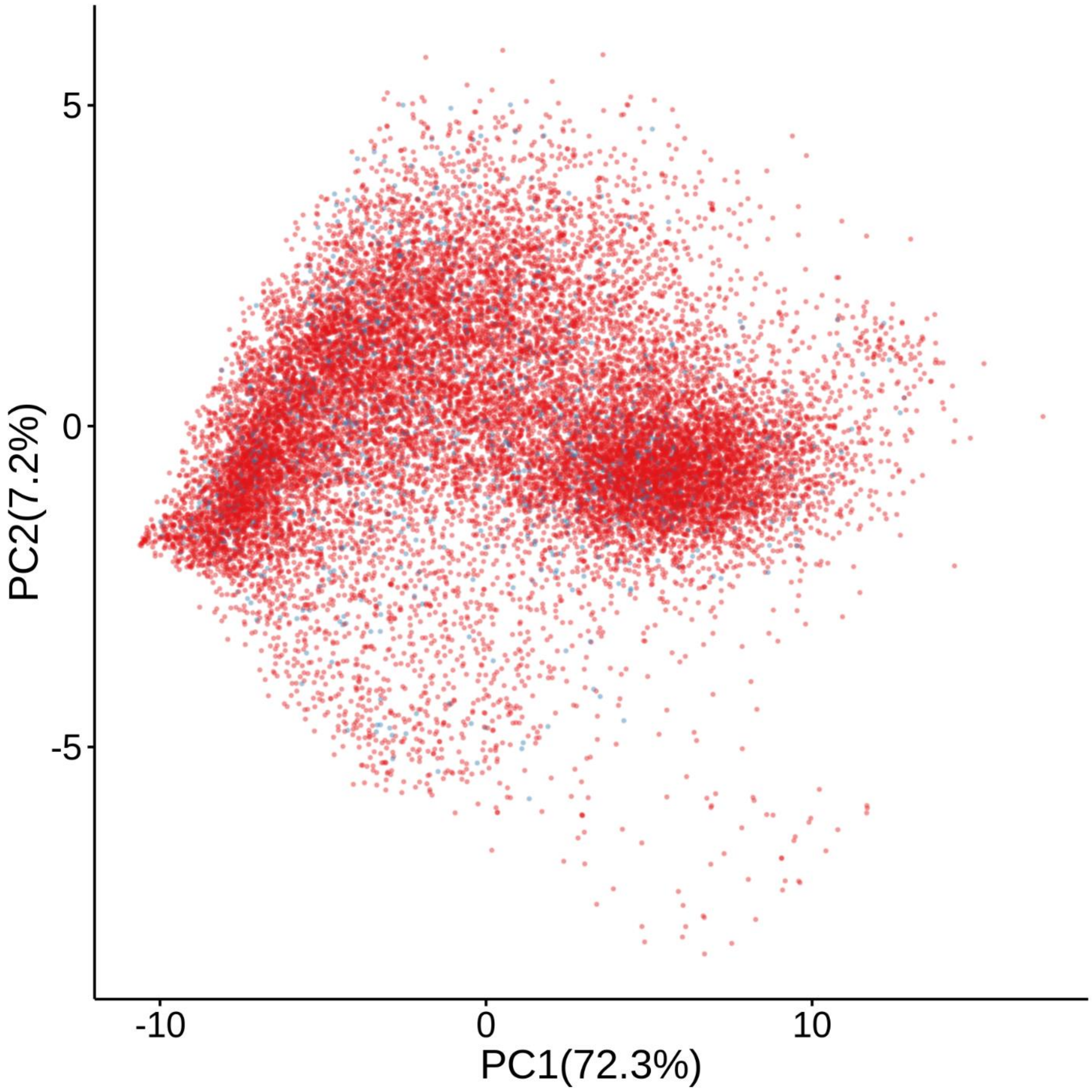


Supplemental Figure 27. Principal components analysis of transcriptomic and WormBase data. Points colored by loss when considering the *Elegans* group. Lost genes do not appear to cluster in multidimensional space. Blue, lost genes. Red, retained genes.

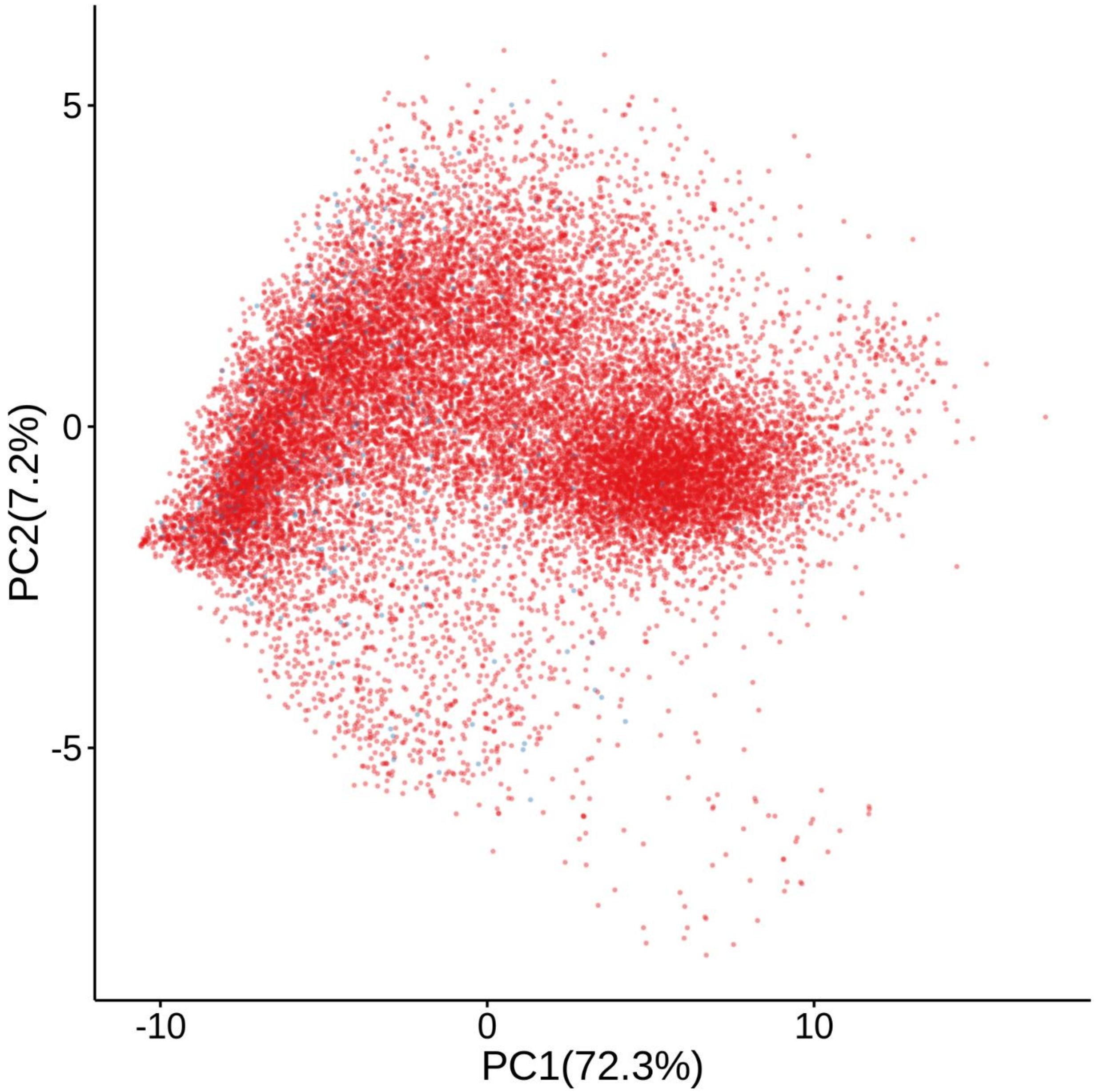


Supplemental Figure 28. Principal components analysis of transcriptomic and WormBase data. Points colored by loss only in *C. inopinata*. Lost genes do not appear to cluster in multidimensional space. Blue, lost genes. Red, retained genes.

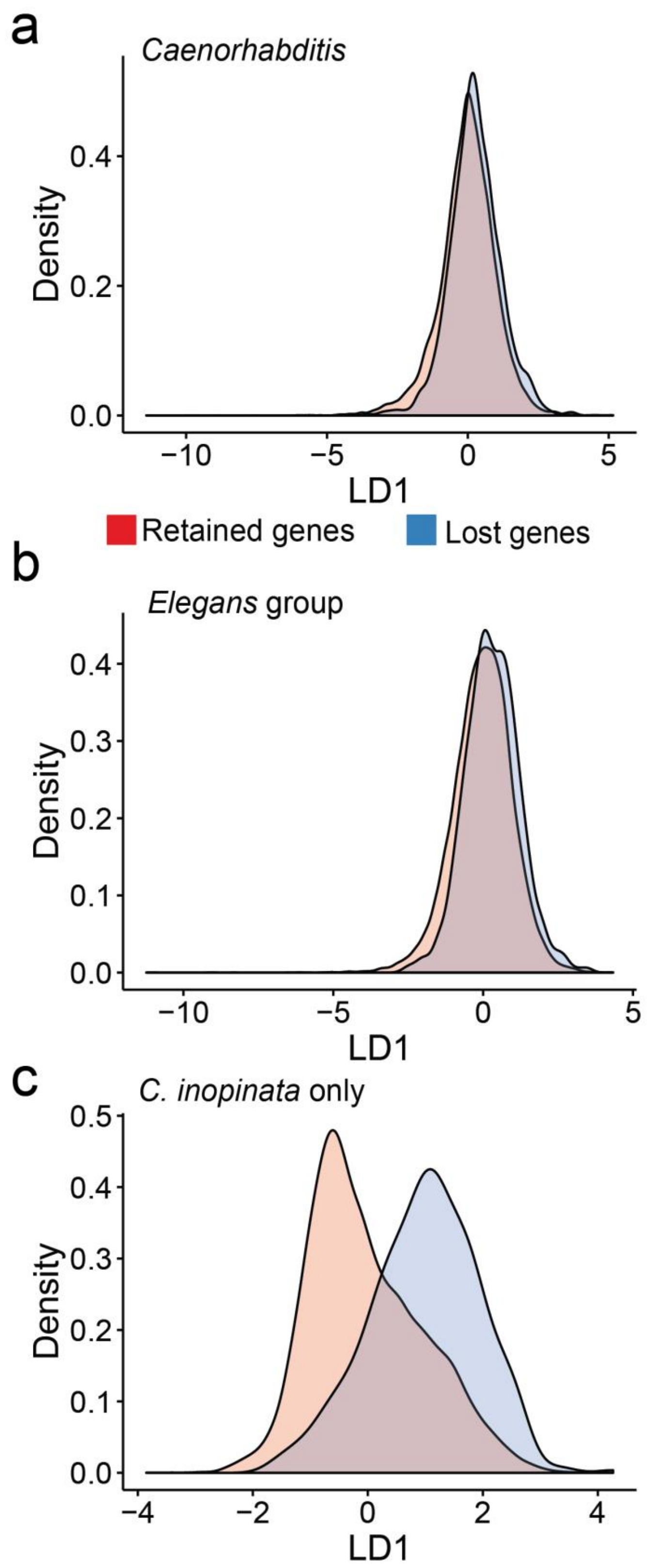


Supplemental Figure 29. Linear discriminant analysis of transcriptomic and WormBase data. Analysis of Boeck et al. 2016 expression data, WormBase ontology term count, and Pfam domain count with genes coded by loss and retained among *Caenorhabditis* species at three phylogenetic levels (all *Caenorhabditis*, a; *Elegans* group, b; and *C. inopinata* only, c). Plotted are the distributions of projected linear discriminants for both gene classes (retained genes and lost genes) across the three levels of phylogenetic consideration.

| **Phenotype** | **Number of genes** |
|---|---:|
| embryonic lethal | 3259 |
| sterile | 2843 |
| lethal | 2034 |
| slow growth | 1964 |
| larval arrest | 1678 |
| locomotion variant | 1531 |
| transgene subcellular localization variant | 1268 |
| reduced brood size | 1189 |
| transgene expression increased | 1113 |
| pattern of transgene expression variant | 1004 |
| maternal sterile | 1002 |
| larval lethal | 861 |
| protruding vulva | 844 |
| transgene expression reduced | 830 |
| sterile progeny | 757 |
| receptor mediated endocytosis defective | 739 |
| dauer lifespan extended | 706 |
| sick | 669 |
| organism development variant | 527 |
| protein aggregation variant | 513 |
| extended life span | 507 |
| avoids bacterial lawn | 482 |
| shortened life span | 461 |
| fat content reduced | 451 |
| germ cell compartment expansion variant | 436 |

Supplemental Table 1. The top 25 most common phenotypes in WormBase.

| **Gene name** | **Sequence name** | **Number of phenotypes** |
|---|---|---|
| *pop-1* | W10C8.2 | 143 |
| *daf-2* | Y55D5A.5 | 134 |
| *daf-16* | R13H8.1 | 98 |
| *glp-1* | F02A9.6 | 87 |
| *mpk-1* | F43C1.2 | 85 |
| *gld-1* | T23G11.3 | 84 |
| *egl-18* | F55A8.1 | 84 |
| *skn-1* | T19E7.2 | 79 |
| *dyn-1* | C02C6.1 | 79 |
| *unc-73* | F55C7.7 | 78 |

Supplemental Table 2. The top ten “most pleiotropic” genes as measured by number of unique WormBase phenotypes.

| **Gene name** | **Sequence name** | **Number of references** |
|---|---|---|
| *daf-16* | R13H8.1 | 760 |
| *daf-2* | Y55D5A.5 | 654 |
| *glp-1* | F02A9.6 | 312 |
| *skn-1* | T19E7.2 | 278 |
| *ced-3* | C48D1.2 | 265 |
| *lin-12* | R107.8 | 240 |
| *age-1* | B0334.8 | 238 |
| *let-60* | ZK792.6 | 224 |
| *unc-54* | F11C3.3 | 195 |
| *daf-12* | F11A1.3 | 188 |

Supplemental Table 3. The top ten “most widely studied” genes as measured by WormBase reference count.
